# Multi-Objective Engineering of Fibrin-Selective Thrombolytic Proteases with Enhanced Biocatalytic Efficiency and Inhibition Resistance

**DOI:** 10.64898/2026.01.19.700241

**Authors:** M. Toul, V. Slonkova, J. Mican, S. Thalerova, M. Peskova, P. Kittova, P. Scheer, J. Hlozkova, E. Brhelova, D.A. Aksu, J. Biskupic, M. Kuchynka, J. Ondrus, P. Kasparek, T. Batkova, M. Marek, J. Vitecek, L. Kubala, R. Mikulik, J. Damborsky, D. Bednar, Z. Prokop

**Author notes:** These authors contributed equally to the work (M. Toul, V. Slonkova, J. Mican).

## Abstract

Thrombolytic enzymes represent an important class of proteolytic biocatalysts for medical applications, yet currently used FDA-approved variants, including alteplase and tenecteplase, remain limited by suboptimal catalytic efficiency, off-target activity, and susceptibility to inhibition. These limitations reflect the complexity of enzyme function in physiological environments, where therapeutic performance depends on the simultaneous optimization of multiple catalytic and biophysical properties. Here, we introduce a multi-objective enzyme engineering strategy for the design of next-generation thrombolytic proteases, explicitly targeting multiple properties required for therapeutic performance. Our approach combines computer-aided design, evolutionary reconstruction, and literature-guided mutation selection to improve catalytic activity, fibrin selectivity, inhibition resistance, and functional lifetime within a single workflow. This framework is coupled with systematic biochemical characterization, in vitro evaluation of clot penetration and fibrinolytic activity, and in vivo validation of efficacy and safety. By addressing multiple performance parameters simultaneously, this strategy enables efficient navigation of trade-offs that typically limit enzyme optimization. Using this approach, we identify Brnoteplase as a lead variant with enhanced fibrin selectivity, improved resistance to inhibition, and superior clot penetration, resulting in increased effective catalytic lifetime and enabling bolus administration. In vivo studies demonstrate enhanced thrombolysis and recanalization with reduced hemorrhagic complications. These findings provide a broadly applicable framework for designing proteolytic biocatalysts suitable for complex biological environments.

## Introduction

Cardiovascular diseases, including ischemic stroke, remain the leading cause of morbidity and mortality worldwide, with a steadily increasing global burden^1^. Although mechanical thrombectomy has significantly improved outcomes in selected patients, its use is inherently limited by the need for specialized infrastructure and narrow treatment windows^2^. In contrast, enzyme-mediated thrombolysis is widely accessible, cost-effective, and straightforward to administer, making it the primary therapeutic option on a global scale^3^. However, currently available thrombolytic enzymes exhibit substantial limitations^4^, underscoring the need for engineering next-generation biocatalysts with improved efficiency and safety profiles^5^.

For nearly three decades, alteplase was the only U.S. Food and Drug Administration (FDA)-approved thrombolytic for acute ischemic stroke. Despite its clinical utility, alteplase shows modest recanalization rates (<50%), a short biological half-life, and an elevated risk of intracranial hemorrhage^6,7^. The recent FDA approval of tenecteplase in March 2025 introduced a second drug for acute ischemic stroke treatment^8^. As an engineered variant of alteplase, tenecteplase exhibits improved fibrin specificity and improved inhibition resistance, resulting in an extended half-life that enables bolus administration^9^. However, tenecteplase does not show clear superiority over alteplase in terms of efficacy or safety^10–13^, highlighting persistent limitations of current thrombolytic enzymes.

From a catalytic perspective, thrombolytic performance arises from the interplay of multiple enzyme properties. An ideal thrombolytic enzyme should combine high catalytic efficiency toward fibrin-bound plasminogen with strict substrate selectivity and reduced binding to receptors such as the low-density lipoprotein receptor-related protein 1 (LRP1) and the N-methyl-D-aspartate receptor (NMDAR)^14^. A combination of these properties would make the 69 thrombolytic safer by decreasing hemorrhagic and neurotoxicity risks^15,16^. In parallel, enhanced resistance to plasminogen activator inhibitor-1 (PAI-1) prolongs the biological half-life, thereby increasing overall efficacy throughout the therapeutic window and enabling convenient bolus administration^17^. These requirements define a multidimensional optimization problem in which improving a single parameter is insufficient to achieve overall therapeutic performance.

Here, we introduce a multi-objective enzyme engineering strategy to develop next-generation thrombolytic proteases with enhanced biocatalytic and therapeutic properties (**Figure 1**). The workflow integrates complementary design principles, including computer-aided mutagenesis to modulate receptor interactions, literature-guided combination of beneficial mutations, ancestral sequence reconstruction to access robust sequence variants, and sequence database mining to identify novel functional scaffolds. These design strategies are combined with systematic biochemical profiling, and hierarchical functional screening to efficiently navigate sequence space while minimizing reliance on resource-intensive *in vivo* testing.

**Figure 1:**
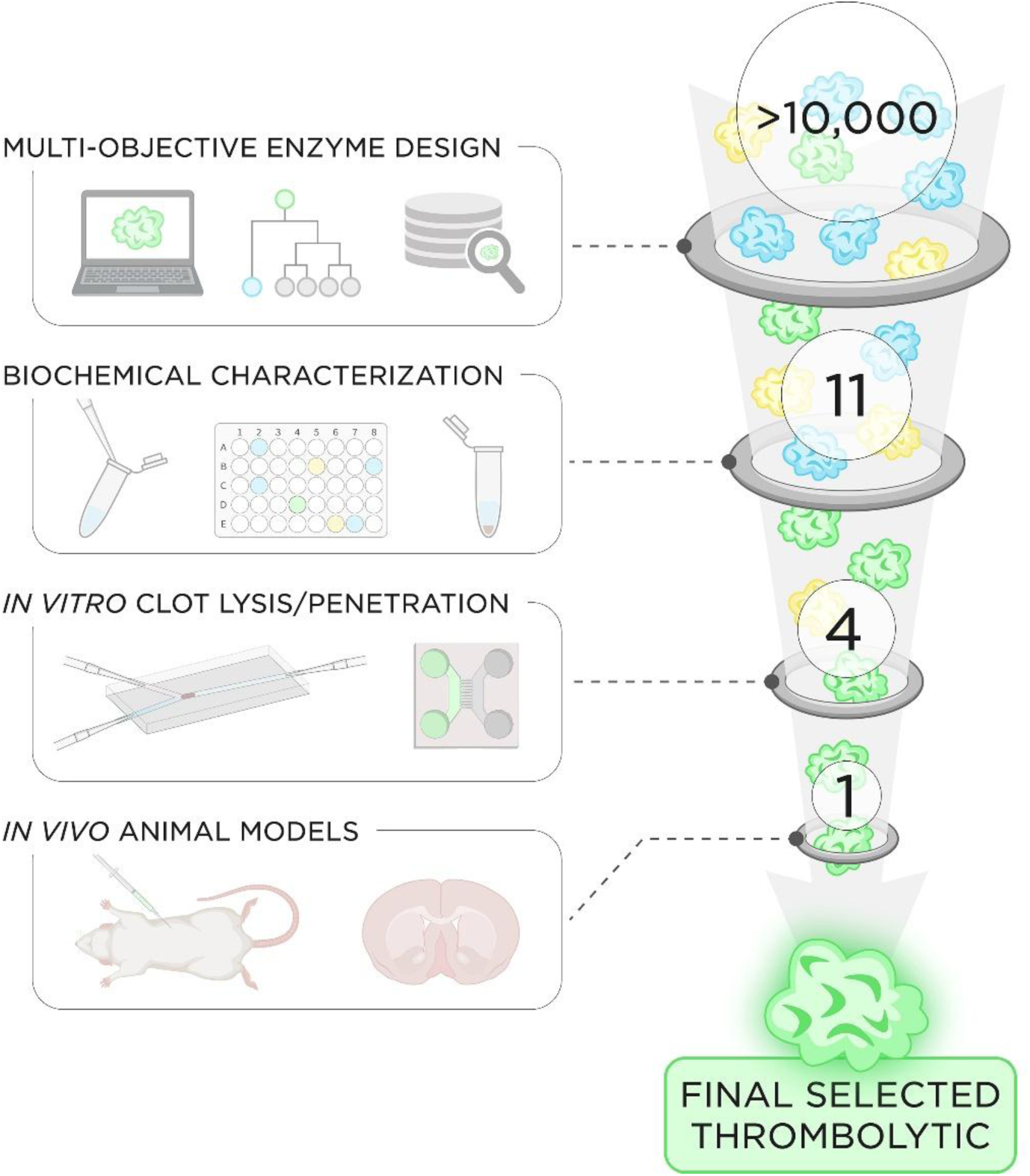
Multi-objective engineering workflow for the development of thrombolytic proteases. The pipeline involves (i) *in silico* rational design of new thrombolytic protein sequences, (ii) their biochemical characterization, (iii) testing in *in vitro* thrombolysis and penetration assays, and (iv) validation in *in vivo* animal rat models. The numbers in the circles correspond to the number of thrombolytic protein variants advanced to each step (grey rings) of the workflow, enabling initial screening of more than 10,000 variants and their comprehensive filtering down to a single selected thrombolytic variant for verification in resource-intensive preclinical *in vivo* models.

Using this approach, we engineered variants of plasminogen activators and identified a lead candidate, Brnoteplase, with enhanced fibrin selectivity, resistance to inhibition, and improved clot penetration. These improvements translate into prolonged effective catalytic lifetime and improved thrombolytic performance, positioning Brnoteplase as a promising candidate for further development. Beyond this specific case, our results establish a generalizable framework for the multi-objective engineering of proteolytic biocatalysts operating in complex biological environments, laying a foundation for the development of next-generation enzyme-based therapeutics.

## Methods

### Computer-aided selection of mutations alleviating binding of tPA to LRP1

Computational design of alteplase variants with reduced binding to the LRP1 receptor was performed using (i) structure-based and (ii) sequence-based strategies.

In the structure-based strategy, we performed conservation and correlation analysis of lysine residues on the first four domains of alteplase, which are responsible for LRP1 binding, using the HotSpot Wizard web server^18^. Afterwards, we docked the structure of LRP1 complement-like binding repeat to tissue-type plasminogen activator (tPA) using ClusPro and RosettaDock^19^. We clustered the docked structures and evaluated distances of conserved and exposed lysines of tPA to the lysine-binding site of LRP1. Lastly, we have performed molecular dynamics (MD) of tPA and evaluated which pairs of lysines are in the required distance of 21 to 29 Å for binding LRP1^20^. We designed mutant tPAs lacking specific lysines. These lysines fulfil the following criteria: (i) conserved in homologous sequences, (ii) exposed on the protein surface, (iii) located on the first four domains of tPA responsible for LRP1 binding, (iv) located on the interface with LRP1 complement-binding repeat in cluster centres of docked structures, and (v) being within 21 – 29 Å of another lysine residue in tPA during MD simulations.

In the sequence-based strategy, we compared the tPA sequence to its homolog, desmoteplase, which competitively binds LRP1^21^. In the pairwise sequence alignment, we identified lysines in both tPA and desmoteplase at homologous positions. These lysines are likely candidates to mediate LRP1 binding.

Lysine positions identified by structure– and sequence-based strategies were subjected to *in silico* saturation mutagenesis using the Rosetta ddg_monomer application. Mutated residues that retained thermodynamical stability were selected for experimental characterization.

### Combining beneficial tPA mutations from literature

To identify mutations for enhancing the efficacy and safety of tPA, we performed a comprehensive literature review on the structural biology and protein engineering of thrombolytics. We then sought an innovative combination of mutations to improve tPA efficacy and safety and combined them into a multiple mutant.

### Ancestral sequence reconstruction of tPA-related proteins

The sequence of tPA, including the signal peptide, was submitted to the FireProt^ASR^ web server^22^. The calculation was performed using default settings. Ancestral sequences for experimental characterization were selected using the following criteria: (i) as few insertions or deletions compared to tPA as possible, (ii) preserved catalytic residues, and (iii) preserved functionally important residues known to influence different tPA functions.

### Mining novel enzymes from sequence databases

EnzymeMiner was used to search for novel thrombolytic enzymes in protein sequence databases^23^. Two separate runs were performed using tPA and desmoteplase as query sequences. Proteins with transmembrane regions or mutated catalytic residues were discarded. Moreover, proteins had to contain key domains for plasminogen activation and fibrin binding and score a predicted solubility higher than the query sequence. After filtering for these characteristics, proteins with interesting properties were selected manually.

### Construction of mutant gene sequences

For each designed thrombolytic variant, a synthetic gene (Genscript) was cloned into the expression vector pcDNA3.1 (Thermo Fisher Scientific). *E. coli* strain TOP10F (Thermo Fisher Scientific) was transformed with the plasmid construct. Plasmid DNA was isolated for transient transfection using Plasmid Giga Kit (Qiagen).

### Heterologous gene expression and protein isolation

HEK-293E cells (ATCC® CRL-10852™) were adapted to suspension growth in HyClone™ CDM4HEK293 serum-free medium (Cytiva) supplemented with 4 mM L-glutamine (AppliChem). Prior to transient transfection, HEK-293E cells were centrifuged at 250 g for 10 minutes and resuspended at the density of 2×10^7^ cells/ml in the Gibco™ FreeStyle™ 293 expression medium (Thermo Fisher Scientific). HEK-293E cells were transiently transfected by plasmid DNA and linear 25-kDa polyethyleneimine (Polysciences) at a 1:2 ratio and a final concentration of 25 and 50 ug/mL, respectively. Plasmid pmaxGFP at 5 % of the total DNA content was added as a positive control of the transfection process. Four hours after transfection, cells were diluted with HyClone medium containing 2 mM L-glutamine. The cells were cultured at 37 °C and 130 rpm in an incubator shaker (New Brunswick Innova® 44/44R). The culture supernatants were harvested 7 days post-transfection. The supernatant was purified on Ni-NTA column (Protino, Macherey-Nagel) connected to the BioLogic DuoFlow FPLC system (Bio-Rad) and equilibrated with the purification buffer (10 mM Tris-HCl pH 7.5, 200 mM NaCl). The thrombolytic protein bound to the column was washed and eluted with 10 mM and 250 mM imidazole in buffer 10 mM Tris-HCl buffer pH 7.5 with 200 mM NaCl and 200 mM L-Arginine. Subsequently, the protein was purified on the Superose 6 SEC column (GE Healthcare) equilibrated with PBS buffer pH 7.4, 200 mM L-Arginine. The protein was supplemented with 0.01 % of Tween 80 and stored at +4 °C.

### Protein thermostability measurement

Protein thermostability was determined by analyzing changes in protein intrinsic tryptophan fluorescence with increasing temperature using the nanoscale differential scanning fluorimetry (nanoDSF) instrument Prometheus NT.48 (NanoTemper Technologies). Each measurement was performed in triplicate in phosphate-buffered saline PBS pH 7.4 (10 mM Na_2_HPO_4_, 1.8 mM KH_2_PO_4_, 2.7 mM KCl, 137 mM NaCl) containing 3.5 % L-arginine, and 0.01 % Tween 80. The fluorescence was monitored at temperatures ranging from 20 °C to 100 °C with a temperature ramp of 1 °C.min^−1^ and the values of the fluorescence ratio at 350/330 nm were plotted against temperature. The values of onset temperature (*T*_onset_) and melting temperature (*T*_m_) were determined using the manufacturer-provided software after deriving the obtained fluorescence curves. The final values were expressed as averages of individual replicates and their respective standard deviations.

### Determination of enzymatic activity and fibrin(ogen) stimulation and selectivity

The activities of thrombolytic enzymes were measured using the fluorogenic substrate D-VLK-AMC (AAT Bioquest), which is converted by the generated plasmin molecules to the fluorescent product 7-amino-4-methylcoumarin (AMC), whose fluorescence was monitored based on the previously published protocol^24^. All the experiments were performed at 37 °C in phosphate-buffered saline PBS pH 7.4 (10 mM Na_2_HPO_4_, 1.8 mM KH_2_PO_4_, 2.7 mM KCl, 137 mM NaCl) containing 1 mM CaCl_2_, 0.0035 % L-arginine, and 0.01 % Tween 80 and each measurement was repeated in three replicates. The activities were determined in the absence but also in the presence of fibrinogen or fibrin that served as stimulation agents. In the case of fibrin stimulation, small fibrin clots were formed in a microplate well of a black clear bottom 96-well plate by mixing 8 µL of human fibrinogen (HYPHEN BioMed) with 2 µL of human thrombin (Sigma-Aldrich). The wells were sealed with an adhesive foil to prevent evaporation and kept at room temperature for an hour to ensure complete polymerization and clot formation. In the case of fibrinogen stimulation, the microplate wells were filled with 8 µL of human fibrinogen mixed with 2 µL of the reaction buffer only. In the absence of any stimulant, only 10 µL of the reaction buffer was injected. After that, all the microplate wells were added 50 µL of human plasminogen (Roche Diagnostics), 30 µL of D-VLK-AMC, and 10 µL of the tested thrombolytic to initiate the reaction. The resulting concentrations in microplate wells were 3.0 µM fibrin (where applicable), 0.04 UN/mL thrombin (where applicable), 3.0 µM fibrinogen (where applicable), 4.5 µM plasminogen, 200 µM D-VLK-AMC, and 0.08 nM tested thrombolytic. The microplate containing all the tested conditions was then sealed with a transparent adhesive foil and the enzymatic activity was measured by monitoring the increase in fluorescence with time using the microplate reader FLUOstar OPTIMA (BMG Labtech). The emission was detected every 60 seconds at 460 nm upon excitation at 360 nm with the instrument gain set to 1,100 and the bottom reading position.

### Activity and stimulation kinetic data analysis

All the curves of increasing AMC fluorescence with time were first recalculated to AMC concentration changes with time based on the calibration curve that was obtained by measuring fluorescence of various increasing concentrations of the AMC standard (Sigma-Aldrich) ranging from 0 to 1000 µM. The concentration dependence exhibited an exponential trend, so the calibration curve was constructed by fitting the calibration points with the exponential **Equation 1** to obtain the calibration factors *F*_a_, *F*_b_, and *F*_c_.

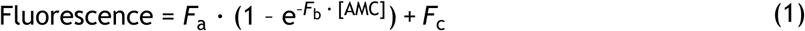

Due to the coupled nature of the applied assay, the initial phase of each recalculated kinetic curve was fitted with the quadratic **Equation 2**, where *v*_0_ corresponds to the determined initial rate of the enzymatic reaction, *t*_0_ to the dead-time between the manual mixing of the assay components and the start of the measurement, and *c* to the initial y-axis offset. Final values of initial rates (activities) were expressed as average ± standard deviation for each measured sample and condition.

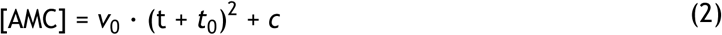

Stimulation factors and selectivities were determined according to the previously postulated definitions^25^. Stimulation factor for the fibrinogen stimulation (*Stim*_FG_) or the fibrin stimulation (*Stim*_FN_) was calculated as a ratio of the activity in the presence of the stimulant (*v*_0,FG_ or *v*_0,FN_) over the activity in the absence of the stimulant (*v*_0_), as summarized in **Equation 3 and 4**. Selectivity (*Sel*_FN_) towards fibrin (with respect to fibrinogen) was then calculated as a ratio of stimulation factors in the presence of fibrin over fibrinogen according to **Equation 5**. The resulting value was relativized to alteplase and statistically compared using two-sample *t*-test.

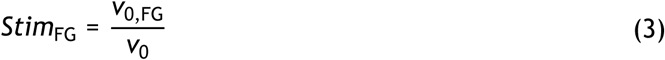

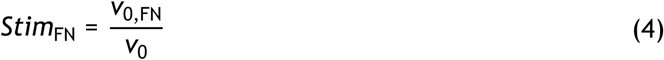

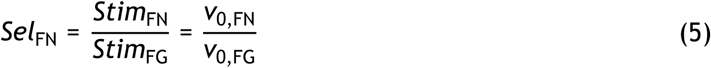

### Plasminogen activator inhibitor 1 (PAI-1) inhibition resistance

The potency of thrombolytic inhibition was analyzed by determining the fraction of active thrombolytics at different PAI-1 concentrations based on the previously published protocol^24^. The experiment was performed at 37 °C in phosphate-buffered saline PBS pH 7.4 (10 mM Na_2_HPO_4_, 1.8 mM KH_2_PO_4_, 2.7 mM KCl, 137 mM NaCl) containing 1 mM CaCl_2_, 0.0035 % L-arginine, and 0.01 % Tween 80, and each measurement was repeated in three replicates. 40 µL of PAI-1 (Sigma-Aldrich) was mixed with 10 µL of a tested thrombolytic and the sealed mixture was incubated at 37 °C for 60 min while shaking at 180 rpm to ensure that the interaction with PAI-1 reached equilibrium. The resulting concentrations in the inhibition mixture were 0–1.5 µM PAI-1 and 0.16 µM thrombolytic. Residual activity was determined by transferring 10 µL of the inhibition mixture into a microplate well of a black clear bottomed 96-well plate and mixing it with 40 µL of the tPA fluorogenic substrate Boc-L-(p-F)FPR-ANSN-H-C_2_H_5_ (US Biological) to initiate the enzymatic reaction. The resulting fluorogenic substrate concentration was 200 µM. The microplate was sealed with a transparent adhesive foil and the enzymatic activity was detected by monitoring the increase in fluorescence over time using the microplate reader FLUOstar Omega (BMG Labtech). The emission was detected every 60 seconds at 460 nm upon excitation at 360 nm with the instrument gain set to 580 and the bottom reading position.

### Inhibition resistance data analysis

The linear phase of each fluorescence kinetic curve was fitted with the linear **Equation 6** where the slope *v*_i_ corresponds to the rate of the inhibited enzymatic reaction (residual activity) and *F*_0_ to the initial fluorescence offset. Since the fluorogenic substrate was converted directly by the tested thrombolytic enzymes with no reaction coupling and since the fluorescence of the generated product was directly proportional to its concentration, fluorescence curves could be fitted directly with the linear equation to reliably obtain relative values of initial rates without a need for recalculation to concentration changes. Final values of initial rates were expressed as average ± standard deviation for each measured sample and condition.

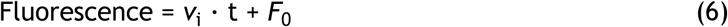

Concentration dependence of determined residual activities *v*_i_ with increasing PAI-1 concentration was fit with **Equation 7** for a tightly bound inhibitor where c_I_ corresponds to the total concentration of PAI-1, c_E_ to the total concentration of a thrombolytic enzyme, *v*_i_ to the residual activity at a given concentration c_I_ of PAI-1, *v*_0_ to the non-inhibited activity in the absence of PAI-1, and *IC*_50_ to the concentration of PAI-1 causing 50 % inhibition of a tested thrombolytic. The values of *IC*_50_ were determined for each tested thrombolytic, relativized to the value for alteplase, and statistically compared using a two-sample *t*-test.

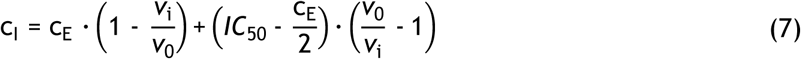

### Fibrin binding and affinity

Affinities of thrombolytics towards fibrin were analyzed by indirectly determining the level of fibrin-bound thrombolytics at various fibrin concentrations based on the previously published protocol^24^. The experiment was performed at 37 °C in phosphate-buffed saline PBS pH 7.4 (10 mM Na_2_HPO_4_, 1.8 mM KH_2_PO_4_, 2.7 mM KCl, 137 mM NaCl) containing 1 mM CaCl_2_, 0.0035 % L-arginine, and 0.01 % Tween 80 and each measurement was repeated in three replicates. 320 µL of plasminogen-free human fibrinogen (HYPHEN BioMed) was premixed with 40 µL of a tested thrombolytic and the clotting was initiated by the addition of 40 µL of human thrombin (Sigma-Aldrich). The resulting concentrations in the polymerization mixture were 0–38 µM fibrinogen, 16 nM (alteplase, Alt01, Alt02, Alt03, AltAnc02, AltEM03, DesEM01) or 235 nM (tenecteplase, Brnoteplase, DesEM02, desmoteplase) thrombolytic, and 2 UN/mL thrombin. This mixture was incubated at 37 °C for 60 min while shaking at 180 rpm to ensure that a complete clot formation was achieved. The clots with bound thrombolytics were separated by centrifugation at 4 °C and 15,000 rpm for 30 minutes and the amount of unbound thrombolytic in the supernatant was determined by residual activity measurement. 20 µL of the supernatant was carefully transferred into a microplate well of a black clear bottom 96-well plate and mixed with 50 µL of human plasminogen (Roche Diagnostics) and 30 µL of the fluorogenic substrate D-VLK-AMC (AAT Bioquest) to initiate the enzymatic reaction. The resulting concentrations in a microplate well were 2.3 µM plasminogen and 200 µM D-VLK-AMC. The microplate was sealed with a transparent adhesive foil and the enzymatic activity was detected by monitoring an increase of fluorescence with time using the microplate reader FLUOstar OPTIMA (BMG Labtech). The emission was detected every 60 seconds at 460 nm upon excitation at 360 nm with the instrument gain set to 1,100 and the bottom reading position.

### Fibrin binding equilibrium data analysis

The values of residual plasminogen activation initial rates (activities) *v*_0,res_, corresponding to the relative amount of an unbound thrombolytic agent, were determined by fitting the kinetic curves as described above in the corresponding ‘Activity and stimulation kinetic data analysis’ section. These values were then relativized to the initial rate in the absence of fibrin (*v*_0,abs_) according to **Equation 8** in order to obtain a fraction of bound plasminogen activator (*Bind*) at a given concentration of fibrin. Concentration dependence of the determined bound fraction *Bind* with increasing fibrin concentration was fit with the hyperbolic **Equation 9** to obtain the value of dissociation constant *K*_d_ and the percentage of the maximal fibrin binding *Bind*_lim_. The values were determined for each tested thrombolytic, relativized to the value for alteplase, and statistically compared using a two-sample *t*-test.

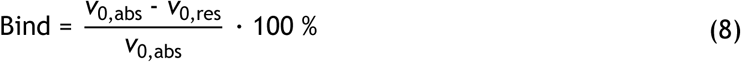

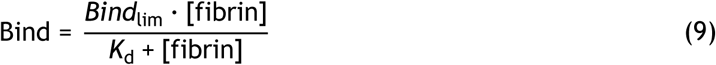

### Preparation of clots and fibrin gel for *in vitro* assays

Semi-synthetic clots were prepared by mixing human fibrinogen (Tisseel Lyo kit, Baxter, dissolved in water for injection to reach the concentration of 52 mg/mL), human thrombin (Tisseel Lyo kit, Baxter, dissolved in 40 µM CaCl_2_ solution to the concentration of 3.3 IU/mL), and red blood cells (RBCs) isolated from healthy donors’ blood in the volume ratio of 1:1:1. The incubation and clotting were performed in plastic Eppendorf tubes (0.5 mL for static model; 0.2 mL for flow model) for 30 min at 37 °C and >90 % relative humidity under static conditions. The prepared clots were immediately introduced into the model. The final concentrations of components in the prepared clots were 17.3 mg/mL fibrinogen, 1.1 IU/mL thrombin, and 4.2·10^9^ RBCs/mL and the total clot volume was 90 µL for the static model and 60 µL for the flow model.

RBC dominant clots were prepared from 200 µL of healthy donors’ whole blood without anticoagulant by clotting in borax glass tubes (inner diameter 8 mm) for 3.5 h at 37 °C and >90 % relative humidity under static conditions. The prepared clots were immediately introduced into the model.

Fibrin gel was prepared from the mixture of fibrinogen (Tisseel Lyo kit, Baxter, dissolved in water for injection to the concentration of 34.6 mg/mL) and thrombin (Tisseel Lyo kit, Baxter, dissolved in 40 µM CaCl_2_ solution to the concentration of 2.2 IU/mL) in the volume ratio of 1:1. The incubation and the clotting was performed inside micro-slide channels for 30 min at 37 °C and >90 % relative humidity under static conditions. The final concentrations of the components in the prepared fibrin gel were 17.3 mg/mL fibrinogen and 1.1 IU/mL thrombin and the total volume per micro-slide channel was 30 µL.

All blood donors had agreed to donate blood samples on the premise of signed informed consent for the collection of blood. Individuals, who had received acetylsalicylic acid, non-steroidal anti-inflammatory, or antiplatelet drugs within 7 days before blood collection, were excluded.

### *In vitro* thrombolytic models

Thrombolytic experiments were performed in static and flow *in vitro* models that were described in contemporary literature^26,27^ but were optimized to determine the suitably measurable effect of alteplase in a highly reproducible manner, as reported in previous publications^28–31^. Briefly, the static model consisted of 1.5 mL plastic Eppendorf tubes filled with fresh frozen human plasma (177RM, Monobind Inc.) medium to a total volume of 500 µL, in which the clots were individually incubated. The tubes were incubated at 37 °C in a dry-block incubator for 60 minutes (the same amount of time as indicated for alteplase treatment of stroke patients).

The flow model consisted of silicone chips made of Sylgard 184 Silicone Elastomer (Dow Corning), which were prepared according to human middle cerebral artery anatomy, with narrowings dimensionally based on patient CT scans (n = 4). The bifurcation was included to enable permanent circulation in the system. Each silicone chip was connected by plastic pipes (internal diameter 3.1 mm) to the 8-channel pump head peristaltic pump MINIPULS 3 (Gilson Inc.), and the entire system was maintained at 37 °C for incubation lasting 180 min. This arrangement maintained the hydrodynamic forces involved in clot removal^32^, whereby a pressure gradient of 10 mm Hg was generated across the occlusion.

Both semi-synthetic and red blood cell dominant clots (preparation described in the corresponding ‘Preparation of clots and fibrin gel for in vitro assays’ section above) were separately employed in both thrombolytic models and the thrombolytic efficiencies were compared. The final concentrations of tested thrombolytic enzymes were selected to be consistent with the clinically relevant dosage indicated for patients with ischemic stroke (1.3 µg/mL), according to the manufacturer’s instructions and supporting pharmacokinetic data^33^. Alternatively, 10-fold lower and 2-, 5-, 10-, and 50-fold higher concentrations were tested as well.

### *In vitro* lytic efficacy analysis

Clot lysis in the static model was determined by measuring clot mass loss^27–29,31,34^ and by spectrophotometric determination of RBC release into the incubation medium at 575 nm^28,29,31^. Clot lysis in the flow model was determined by measuring recanalization time, clot length changes, and spectrophotometric determination of RBC release^28,30,31^. Recanalization frequency was determined as a percentage ratio of complete recanalization to the total number of samples in the respective treatment group. Relative clot reduction was determined as a percentage of clot area reduction prior to clot release from the narrowed occlusion site. In addition, the thrombolysis rate was calculated using linear regression of relative clot reduction at 10-minute intervals. Data were expressed as mean ± standard deviation, if not otherwise indicated. The generated box plots were presented as mean value (cross), median (line), interquartile range (box), and minimum and maximum value (whiskers). ANOVA with post-hoc Tukey’s HSD test was used to compare data. P-values ≤ 0.05 were considered to be statistically significant. All analyses were performed using GraphPad Prism (GraphPad Software Inc.).

### *In vitro* penetration microarray

Penetration experiments were performed in the microarray model outlined in recent literature^35–39^ and optimized to provide a suitably measurable effect of alteplase in a highly repeatable manner. The microarray consisted of the microslide µ-Slide VI 0.4 (Ibidi GmbH) containing channels that were filled with a prepared fibrin gel (procedure described in the corresponding ‘Preparation of clots and fibrin gel for in vitro assays’ section above). Fluorescently labelled plasminogen activators or bovine serum albumin (A9418, Sigma-Aldrich), as a relevant fibrin non-interacting control, were applied to individual microslide channels and fluorescence intensities were continuously recorded in 5 min intervals for 180 min, at 0.25 mm distance intervals up to 1.00 mm from the application site. The fluorescence signal was recorded using the microscope ZEISS Axio Observer Z1 (Carl Zeiss AG) equipped with the metal halide fluorescence light source HXP 120 (Carl Zeiss AG) and the high efficiency Filter Set 43 HE (Carl Zeiss AG). The detection settings were as follows: excitation BP 550/25; beam splitter FT 570; emission BP 605/70. Thrombolytic proteins and bovine serum albumin control were fluorescently labelled with rhodamine B isothiocyanate (283924, Sigma-Aldrich) prior to the penetration assay according to the standard labeling protocol provided by the manufacturer. The final concentrations of the labelled proteins were the same as for thrombolytic experiments, i.e., 1.3 µg/mL. Thrombolytic efficiencies of labelled plasminogen activators were verified in the static model.

### Penetration rate analysis

Protein penetration rate through the fibrin gel was determined by continuous measurements of fluorescence intensities at a distance of up to 1 mm from the front of the plasminogen activator application site^31^. The penetration rate was expressed as the relative fluorescence intensity (RFI) at a given distance and time. RFI was determined using a custom image analysis script in the Python programming language. Penetration rate data were further normalized to relevant controls. Data were expressed as mean ± standard deviation, if not otherwise indicated. The generated box plots were presented as mean value (cross), median (line), interquartile range (box), and minimum and maximum value (whiskers). ANOVA with post-hoc Tukey’s HSD test was used to compare data. P-values ≤ 0.05 were considered to be statistically significant. All analyses were performed using GraphPad Prism (GraphPad Software Inc.).

### Preparation of clots for *in vivo* assays

Details of the characteristics and nature of artificial clots are the subject of a previous project^40^, and their preparation is therefore described only briefly. Commercial human fibrin glue Tisseel kit (Baxter) was used to generate artificial clots for *in vivo* experiments. Saline solution (1.5 mL) and BaSO_4_ suspension (0.5 mL) (Micropaque, Guerbet) were injected into the original “TISSEEL” ampoule. 1 mL of the original calcium chloride solution and 5 mL of bovine serum were injected into the “Thrombin 500 I.U.” original ampoule. Finally, “TISSEEL” and “Thrombin 500 I.U.” preparations were injected in a 1:1 ratio into a PE60 and PE10 tubes (Braintree Scientific) for the thrombolytic effectivity study and the safety study, respectively, using the original Duploject system (Baxter). Both ends of the cannula containing clot components were hermetically sealed to prevent any evaporation or desiccation, and the mixture was incubated at 37 °C for 120 min, followed by storage at 4–8 °C for additional 24 hours. The resulting artificial clot was cut into 10 mm fragments for the effectivity study and 1 mm fragments for the safety study and the corresponding samples were stored in the saline solution in an Eppendorf tube at −20 °C.

### Thrombolytic effectivity study *in vivo*

Adult male outbred Wistar rats (n=68; body weight 120–180 g) were selected for the *in vivo* lysis experiments and were randomized into groups with different treatment regimens (variable thrombolytic types, doses, and administration routes) as described in **Table 1**.

**Table 1:** Overview of animal experimental groups for *in vivo* effectivity study based on different treatment regimes. CRI = continuous rate infusion.

| Experimental group | Number of animals in the group |
| --- | --- |
| Control group (saline solution) | 10 |
| Alteplase (0.9 mg/kg, CRI) | 16 |
| Tenecteplase (0.25 mg/kg, bolus) | 12 |
| Brnoteplase (0.9 mg/kg, CRI) | 4 |
| Brnoteplase (0.9 mg/kg, bolus) | 12 |
| Brnoteplase (2.5 mg/kg, bolus) | 12 |
| Brnoteplase (5.0 mg/kg, bolus) | 2 |

The exclusion criterion for the experiment was death during the anesthesia or the surgical procedures before the clot delivery. Throughout the thrombolysis experiment, rats were under general anesthesia induced by inhalation of 2.5 % isoflurane in filtered room air as a carrier gas. After the intramuscular application of xylazine and ketamine (5 mg/kg and 35 mg/kg of body weight, respectively), a mixture of urethane (80 mg/mL) and alpha-chloralose (6 mg/mL) at a dose of 8 mL/kg of body weight was applied into the peritoneal cavity. Subsequently, the left common carotid artery (CCA) was dissected, visualized, and cannulated in a retrograde direction with a PE50 (Braintree Scientific) cannula.

Three individual barium-labeled artificial clots were injected via the PE50 catheter cannula into the aortic arch of each tested rat. Micro-fluoroscopy using the C-arm LabScope system (Glenbrook Technology, Inc.) was used to verify the localization of the introduced clots in the branches of the abdominal aorta and the position of the animals was then adjusted to achieve the optimal visibility of the clots with the maximum area of their size and without the shielding of bone structures. The initial image was captured at time zero (before the thrombolytic drug administration). Subsequently, a bolus injection/continuous rate infusion of a tested thrombolytic was initiated. The continuous rate infusion treatment consisted of an initial bolus of 10 % of the total dose volume, followed by an infusion of the remaining 90 % volume for 60 minutes. In the bolus treatment groups, the entire volume of the thrombolytic agent was administered at once. The control group received an equivalent volume of intravenous saline solution. The therapy was monitored for 60 minutes by the C-arm micro-fluoroscopy and radiographs were acquired and stored every 5 minutes.

After the experiment, the animals did not regain consciousness. Euthanasia was performed by overdose of anesthetics (0.7 mL of 2 % xylazine + 0.7 mL of 10 % ketamine administered intraperitoneally). Due to the possibility of cross-reactions between analgesics and tested thrombolytic compounds, analgesics were not used.

### *In vivo* thrombolytic effectivity study outcome analysis

The outcome was measured as a relative area change of each monitored artificial clot, and the lysis rate was then calculated as a linear fit of relative area change over time. All collected radiographs were analyzed using the ImageJ 1.52a software (Wayne Rasband, National Institutes of Health), and the area of identified clots in pixels was manually determined at each time point. All images of one animal were measured simultaneously in one reading sequence. Two independent observers performed the image analysis, blinded to one another’s results and to animal group assignment. Anatomical determination of arteries with clots was performed after all measurements. A standardized protocol was established and followed for non-standard clot positions or movements: (1) If the area of two clots in different arteries overlapped, the size was measured as a single clot. (2) If the lysis of the clot caused its movement but the clot was still visible on the image, the shadow area of the clot was included in the original area of the clot. (3) The image was excluded if the clot moved out of the image.

The change in the artificial clot area (CAC) was calculated from the clot area at a calculated time (a_t_) relative to the clot area at time zero baseline before thrombolytics administration (a_0_), i.e., at the beginning of the experiment, based on **Equation 10**. The lysis rate in %.min^−1^ represents a relative change in the area of the artificial clot (CAC) divided by the minute in which the area was measured (t) based on **Equation 11**. Results of the thrombolysis were used to generate the thrombolytic curve. The area under the thrombolytic curve (AUC) was calculated and used as a comparative parameter among the tested and control groups. All analyses were performed using Excel for Microsoft 365 MSO version 2405 (Microsoft Corporation) and GraphPad Prism version 8.0.0 (GraphPad Software Inc.).

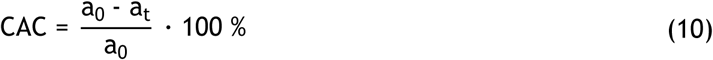

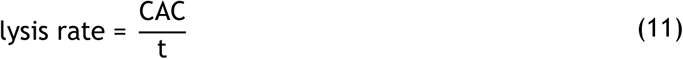

### Safety study

The safety study was based on a thromboembolic Middle Cerebral Artery Occlusion (MCAO) model, utilizing the same clot type as in the thrombolytic effectivity study. However, embolization to all major cerebral arteries, namely the anterior, middle, and posterior cerebral arteries, was included. This approach is referred to as the **C**ircle **o**f **W**illis **B**ranches **O**cclus**i**on – CoWBOi model. Male outbred Wistar rats, weighing between 480 and 980 grams and aged 7 to 9 months, were housed individually under standard conditions, with a natural light/dark cycle and unrestricted access to food and water. Animals were randomly assigned to one of three experimental groups, receiving either alteplase (0.9 mg/kg, continuous rate infusion, n=25, included n=22), tenecteplase (0.25 mg/kg, bolus, n=25, included n=24), or Brnoteplase (2.5 mg/kg, bolus, n=23, included n=22). Rats, that did not develop cerebral stroke or died within 120 minutes of thrombolytic treatment, were excluded from the study.

General anesthesia was induced via inhalation of 2.5 % isoflurane, followed by intramuscular administration of xylazine (5 mg/kg), ketamine (35 mg/kg), and intraperitoneal diazepam (2 mg/kg). Due to the possibility of cross-reactions between analgesics and tested thrombolytic compounds, analgesics were not used. Embolization was performed in a modified version of the standard MCAO model, with cannulation of the left external, and internal carotid arteries. Two embolic clots (1.5 mm × 0.3 mm) were introduced into the intracranial internal carotid artery. Thrombolytic treatment was initiated 4 hours post-embolization. After 24 hours, the animals were euthanized by overdose of anesthetics (0.7 mL of 2 % xylazine + 0.7 mL of 10 % ketamine administered intraperitoneally), and their brains were collected in cold saline without flushing. As described previously, the extracted brains were fixed in formalin for 24 hours and stained with iodine^41^. In contrast with the original set-up, the brains were not examined in agarose but in paraffin oil. Micro-computed tomography (miCT) imaging was performed using a SkyScan1276 CMOS (Bruker) scanner, and the acquired scans were analyzed using the Amira software (Thermo Scientific). Examiners were blinded to group assignments to ensure unbiased assessment.

### *In vivo* safety study outcome analysis

To evaluate hemorrhagic transformation (HT), a classification system based on ECASS II criteria^42^ was applied. Hemorrhagic transformation was categorized into four subtypes (HI1, HI2, PH1, and PH2) according to the criteria listed in **Table 2**. Considering that thrombolytic enzymes were administered 4 hours after clot embolization, by which time all collateral vessels had already closed due to ischemic damage, successful recanalization in this model was consistently associated with hemorrhagic transformation.

**Table 2:** Hemorrhagic transformation (HT) classification based on the ECAS II study. The classification recognizes four HT subtypes (HI1, HI2, PH1, and PH2) that differ in the listed criteria.

| Grade | Description |
| --- | --- |
| HI1 | Petechial hemorrhages at infarct margins |
| HI2 | Petechial hemorrhages throughout the infarct without mass effect |
| PH1 | Hemorrhage affecting $\leq 30$ % of the infarcted area with minor mass effect |
| PH2 | Hemorrhage affecting $> 30$ % of the infarcted area with substantial mass effect |

To quantify hemispheric asymmetry (HA), we calculated the affected hemisphere volume (*affected*) and the intact hemisphere volume (*intact*), excluding the olfactory bulbs and cerebellum from the analysis. The percentage of HA was then derived using **Equation 12**. A hemispheric asymmetry of 6 % was identified as the threshold distinguishing normal from pathological brains. Values exceeding this threshold were consistently associated with the presence of cerebral edema.

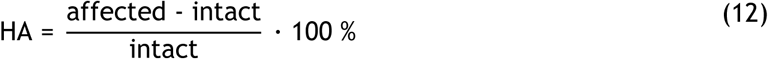

### Quantification of residual plasminogen, plasmin, and thrombolytic levels in rat plasma

At the end of the *in vivo* thrombolytic effectivity study, whole blood from the common carotid was collected to a K3-EDTA tube (2.7 mL, S-Monovette, Sarstedt AG). The collected blood was centrifuged for 15 min at 1,500 rpm and the resulting plasma was separated into an Eppendorf tube, aliquoted, and stored at –20 °C. The collected plasma samples were tested for levels of plasmin, plasminogen, and a thrombolytic enzyme using ELISA Rat Plasmin Kit (MyBioSource), ELISA Rat Plasminogen Kit (MyBioSource), and ELISA Human Active tPA Kit (Molecular Innovations), respectively. The determination of the concentration was performed according to the protocol provided by the manufacturer with no modifications. In the case of Human tPA Kit, purified solutions of alteplase and Brnoteplase (the same as administered in rats) with known concentrations were used to generate individual calibration curves to prevent any discrepancies caused by different sources of the administered and the calibration samples. The final plasma levels were calculated from the measured absorbance values based on the constructed calibration curves.

### Consent for animals used in experiments

All procedures and animal experiments were performed in full compliance with the ARRIVE and the European Community Council Directive (2010/63/EU) for Protection of Vertebrate Animals Used for Experimental and other Scientific Purposes guidelines. The project was approved by the Masaryk University Brno Institutional Committee for the Protection of Experimental Animals and by the Committee for the Protection of Experimental Animals of the Ministry of Education Youth and Sports in the Czech Republic (MEYS CZ) with the protocol number MSMT-35804 / 2019-2.

## Results

### Computational design guides multi-objective selection of thrombolytic candidates

To identify variants with optimized therapeutic performance, we implemented a multi-objective computational design strategy that integrates complementary engineering principles. The initial pool of novel thrombolytic candidates was generated *in silico* using three independent approaches: (i) rational design to modulate active-site interactions and receptor binding, (ii) ancestral sequence reconstruction to access robust and evolutionarily optimized scaffolds, and (iii) database mining to uncover sequence diversity supporting enhanced fibrin selectivity and inhibition resistance (**Table 3**, **Figure S1**). By explicitly considering multiple catalytic and biophysical properties during candidate selection, this framework prioritizes variants with balanced improvements across efficiency, selectivity, functional stability, and safety, providing a focused set of promising enzymes for experimental evaluation.

**Table 3:** Summary of *in silico* designed and selected thrombolytics for experimental testing. Variants highlighted in grey could not be experimentally produced, therefore, were not characterised. Full sequences of all variants are provided in **Figure S1**. ASR = ancestral sequence reconstruction.

| Variant naming | Modification strategy | Parental Protein | Sequence modification |
| --- | --- | --- | --- |
| Alt01 | Rational design | Alteplase | K10E/K49I/Y67N/S69A/K152Q |
| Alt02 | Rational design | Alteplase | Y67N/S69A/K82A/K159S/K162I |
| Alt03 | Rational design | Alteplase | Y67N/S69A |
| Brnoteplase | Rational design | Alteplase | Y67N/S69A/N117Q/W253R/K296A/H297A/R298A/R299A |
| AltAnc01 | ASR | Alteplase | 76.6 % identity to alteplase; ancestral to 5 extant genes |
| AltAnc02 | ASR | Alteplase | 71.0 % identity to alteplase; ancestral to 12 extant genes |
| AltEM01 | Database mining | Alteplase | 78.4 % identity to alteplase; gene isoform X1 from bottlenose dolphin |
| AltEM02 | Database mining | Alteplase | 82.7 % identity to alteplase; gene isoform X2 from bottlenose dolphin |
| AltEM03 | Database mining | Alteplase | 92.0 % identity to alteplase; gene from Bolivian squirrel monkey |
| DesEM01 | Database mining | Desmoteplase | 68.0 % identity to alteplase; gene isoform X3 from Angolan colobus monkey |
| DesEM02 | Database mining | Desmoteplase | 80.0 % identity to alteplase; gene from white-winged vampire bat |

**Rational design** focused on reducing interactions with clearance and signaling receptors, particularly LRP1 and NMDAR, which are implicated in short half-life and neurotoxicity. This was achieved by introducing Tyr67Asn and Ser69Ala mutations shown to prevent glycosylation responsible for LRP1 binding^43^ (resulting in variant Alt03), and by combining these mutations with additional substitutions of lysine residues (yielding variants Alt01 and Alt02). Lysine residues are known to mediate LRP1 interaction. Alt01 includes additional Lys10Glu, Lys49Ile, and Lys152Gln mutations which were selected using structure-based computational tools, namely protein-protein docking, molecular dynamics, surface exposure analysis, and thermodynamic stability predictions (**Supplementary Note 1**). Alt02 incorporates additional Lys82Ala, Lys159Ser, and Lys162Ile mutations selected based on their sequence conservation in desmoteplase, which has been shown to compete with alteplase for the LRP1 binding site^44^ (**Supplementary Note 2**). Finally, the Brnoteplase variant was constructed by combining the two original substitutions preventing LRP1-binding glycosylation^43^ (Tyr67Asn and Ser69Ala) with mutations decreasing PAI-1 inhibition and increasing fibrin specificity^45^ (Lys296Ala, His297Ala, Arg298Ala, Arg299Ala), disabling NMDAR-mediated neurotoxicity^46^ (Trp253Arg), and removing high-mannose glycosylation responsible for the low alteplase biological half-life^47^ (Asn117Gln).

**Ancestral sequence reconstruction (ASR)** strategy utilized the FireProt^ASR^ webserver to explore evolutionary and functional diversity while preserving thrombolytic activity. The analysis produced a phylogenetic tree comprising 150 extant alteplase homologs and 149 ancestral sequences (**Figure S2**). Two ancestral nodes close to alteplase, named AltAnc01 and AltAnc02, were selected based on sequence identity and conservation of catalytic residues (**Table 3**). AltAnc01 is ancestral to alteplase and four other extant sequences annotated as plasminogen activators (PAs), while AltAnc02 is ancestral to alteplase, ten additional annotated PAs, and one hypothetical protein predicted to be a PA (**Supplementary Note 3**).

**Database mining** using the EnzymeMiner webserver identified naturally occurring homologs of alteplase and desmoteplase with potentially favorable biochemical features. Over 10,000 mined sequences were filtered based on predicted solubility, absence of transmembrane regions, and preservation of essential catalytic domains and residues, resulting in 198 alteplase and 615 desmoteplase homologs meeting these criteria. From these, we manually selected five candidates (**Table 3**): three homologous to alteplase (AltEM01–03) and two to desmoteplase (DesEM01–02). The selected sequences originate from bottlenose dolphin, Bolivian squirrel monkey, Angolan colobus, and vampire bat (**Supplementary Note 4**).

### Biochemical profiling reveals Brnoteplase with enhanced thrombolytic properties

Gene sequences encoding putative and mutant plasminogen activators selected *in silico* were experimentally constructed, and 8 out of 11 selected thrombolytic proteins were successfully produced recombinantly. Our comprehensive biochemical characterization (**Figure 2**, **Table S1**) revealed that all the proteins exhibited clear thrombolytic activity (**Figure 2a**, **Figure S3**, **Table S1**), validating the robustness of our computational approach for accurately predicting and designing plasminogen activator enzymes. Thermal stabilities of all characterized proteins (**Figure 2d**, **Figure S4**, **Table S1**) were highly above the physiological temperature, allowing their administration to the human body without a risk of rapid thermal inactivation.

**Figure 2:**
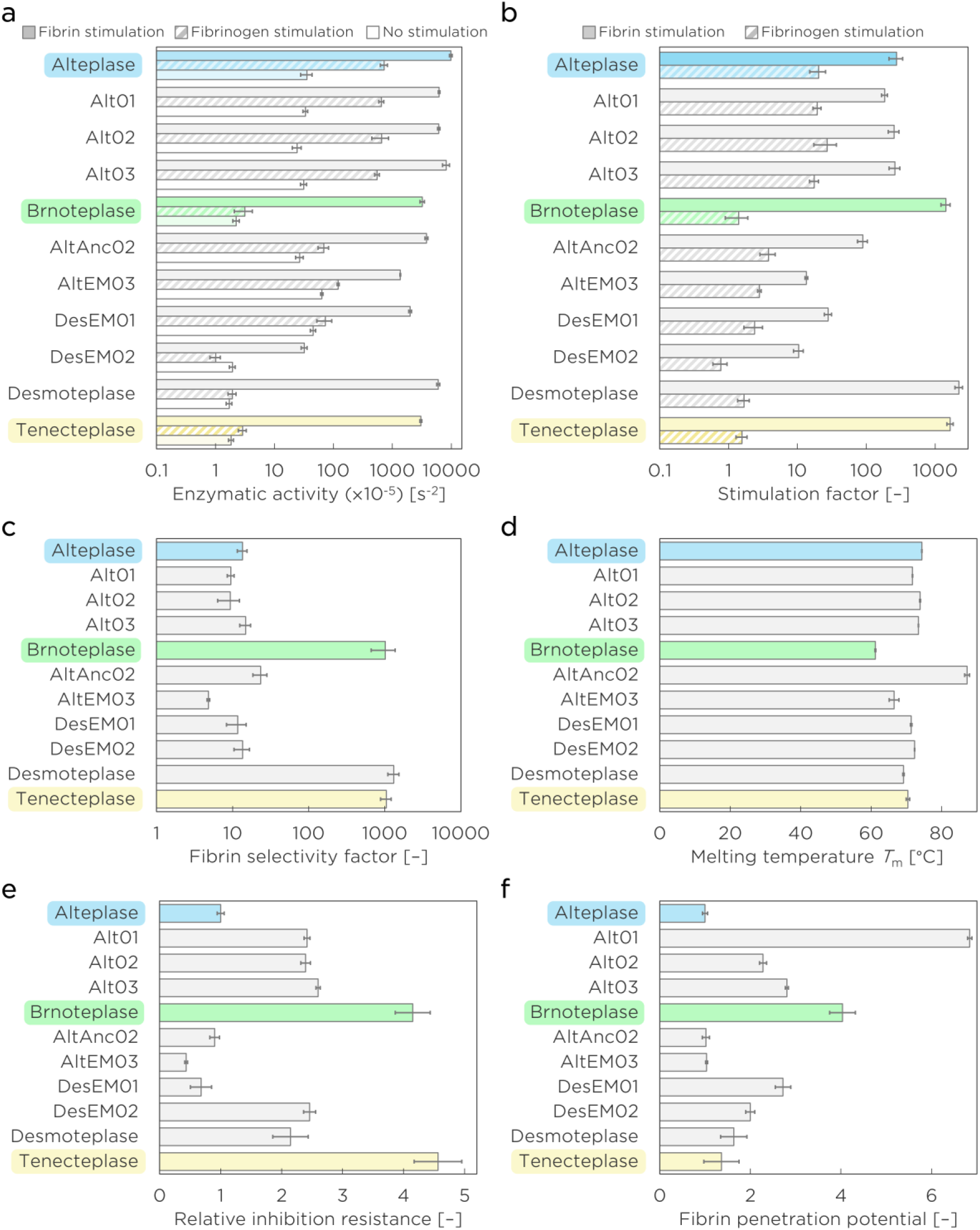
Biochemical characterization of thrombolytic variants. Clinically used variants alteplase and tenecteplase are highlighted in blue and yellow, respectively. The variant Brnoteplase, which was selected for further *in vitro* and *in vivo* testing, is highlighted in green. The bars represent mean values and the whiskers correspond to standard errors. For all measured data and presented values, n=3. (**a**) Comparison of plasminogen activation enzymatic activities (dark solid bars) and their values in the presence of fibrinogen (hatched bars) and fibrin (light solid bars) stimulants. (**b**) Comparison of fibrinogen (hatched bars) and fibrin (solid bars) stimulation factors. The factors are defined as fibrin(ogen)-stimulated activity over non-stimulated activity ratios. (**c**) Comparison of fibrin selectivity factors. (**d**) Comparison of melting temperatures (*T*_m_) to define protein thermal stability. At *T*_m_, half of the protein molecules are unfolded. (**e**) Relative comparison of inhibition resistance towards plasminogen activator inhibitor-1 (PAI-1) calculated as a ratio of the *IC*_50_ value for a tested protein over the *IC*_50_ value for alteplase (reference benchmark protein). (**f**) Relative comparison of fibrin penetration potential calculated as a ratio of the fibrin dissociation constant (*K*_d_) value for a tested protein over the fibrin *K*_d_ value for alteplase (reference benchmark protein). Brnoteplase exhibited superior fibrin stimulation, fibrin selectivity, PAI-1 inhibition resistance, and penetration potential. Please note the logarithmic scale of the x-axis in (**a**), (**b**), and (**c**).

Fibrinogen and fibrin stimulation analysis (**Figure 2b**, **Figure S3**, **Table S1**) yielded unimpaired profiles for Alt01–03 compared to alteplase (p=0.37, p=0.35, p=0.49, respectively), whereas other variants showed significantly lower fibrin stimulation (p<0.001) and activity (p<0.001), suggesting inefficient thrombolysis on fibrin clots. Remarkably, the variant Brnoteplase displayed minimal activity with no stimulation and when stimulated by fibrinogen, but it was massively stimulated by fibrin, 5.2-fold more than alteplase (p<0.001). This resulted in an extraordinary Brnoteplase fibrin selectivity of 1020 ± 360 (**Figure 2c**, **Figure S3**, **Table S1**) that was much higher than for alteplase (13.5 ± 2.0) but similar to highly selective thrombolytics desmoteplase (1320 ± 220) and tenecteplase (1050 ± 170).

All four sufficiently fibrin-active variants, Alt01–03 and Brnoteplase, further exhibited significantly increased resistance to PAI-1 inhibition (p<0.001) and reduced fibrin binding (p<0.001) compared to alteplase, indicating improved potential for clot penetration and lysis throughout the entire volume from inside. The highest inhibition resistance (4.2-fold higher than that of alteplase) was observed for the variant Brnoteplase (**Figure 2e**, **Figures S5 and S6**, **Table S1**), while the most pronounced reduction in fibrin binding was observed for Alt01, followed by Brnoteplase (**Figure 2f**, **Figures S7 and S8**, **Table S1**) by providing 3.9– and 2.3-fold lower affinities compared to that of alteplase, respectively.

In summary, multi-objective biochemical evaluation identified Brnoteplase as a thrombolytic variant with superior functional properties compared to alteplase, combining: (i) 80-fold enhanced fibrin selectivity, (ii) 4-fold increased resistance to PAI-1 inhibition, and (iii) 4-fold improved penetration potential.

### *In vitro* thrombolysis and penetration confirm enhanced Brnoteplase properties

Alt01–03 and Brnoteplase, the four variants identified as highly fibrin-active in biochemical assays, were subjected to thrombolysis testing on *in vitro* prepared human clots and compared to alteplase and tenecteplase (**Figure 3**). Treatment with all tested thrombolytics in the static model containing semi-synthetic clots resulted in significantly greater clot mass loss (p<0.001) and red blood cell (RBC) release (p<0.001) than the untreated group (**Figure 3a**, **Figure S9**, **Table S2**). The results of lysis efficiencies were consistent with the biochemical characterization data (**Figure 2a**).

**Figure 3:**
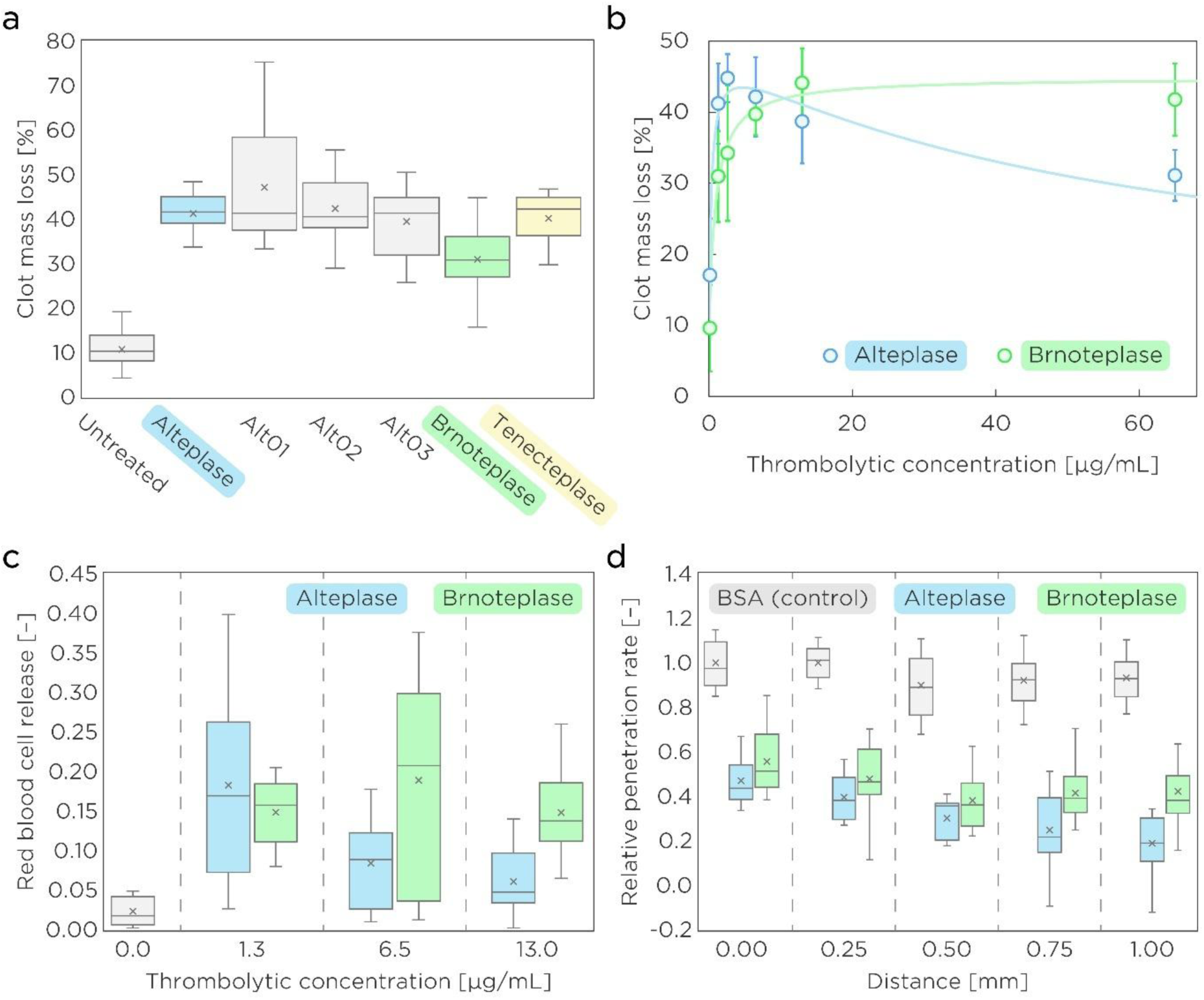
*In vitro* analysis of thrombolysis efficacy and fibrin penetration. (**a**) Comparison of thrombolytic efficacy in the static model with semi-synthetic clots expressed as clot mass loss. (**b**) Concentration dependence profile of thrombolytic efficacy with increasing applied concentration of a tested thrombolytic in the static model. (**c**) Comparison of thrombolytic efficacy at multiple concentrations in the flow model expressed as red blood cell release. (**d**) Comparison of relative penetration rates at multiple distances of a fibrin gel. Box plots illustrate mean values (cross), median (line), interquartile range (box), and minimum/maximum values (whiskers). Scatter plots illustrate mean values (circle) and standard errors (whiskers). The number of independent experiments for each presented value are summarized in **Tables S2–S7**.

The concentration-dependence experiment with alteplase and the most promising variant, Brnoteplase (**Figure 3b**, **Table S3**), revealed notable differences between the two tested proteins. Alteplase showed no significant enhancement of thrombolysis with increasing concentration, as clot mass loss at 2.6, 6.5, and 13 µg/mL was comparable to the clinically relevant concentration of 1.3 µg/mL (p=0.93, p>0.99, p=0.97, respectively) and it even decreased at the highest applied concentration of 65 µg/mL (p=0.002). In contrast, Brnoteplase exhibited a clear trend of increasing thrombolytic efficacy with increasing dosing: clot mass loss was comparable to the clinically relevant concentration only at 2.6 µg/mL (p=0.96) while significantly greater lysis was observed at 6.5, 13, and 65 µg/mL (p=0.02, p<0.001, p<0.001, respectively). At higher concentrations of 6.5 and 13 µg/mL, Brnoteplase reached the comparable level of lysis as for alteplase (p>0.99, p=0.20, respectively), and it even surpassed alteplase efficiency at the highest concentration of 65 µg/mL (p=0.009).

Similar trends were observed for RBC release analysis with semi-synthetic clots (**Figure S10a**, **Table S3**) and RBC-dominant clots (**Figure S10bc**, **Table S4**), confirming the validity of the measured data (details in **Supplementary Note 5**). In the flow model (**Figure 3c**, **Figure S11**, **Table S5**), alteplase provided lower recanalization time (p=0.005), greater clot reduction rate (p<0.001), and greater RBC release (p=0.006) compared to the untreated group only at the clinically relevant concentration (1.3 µg/mL), while higher concentrations yielded no significant effect compared to the untreated group. In contrast, Brnoteplase was significantly more effective than the untreated group across all tested concentrations (details in **Supplementary Note 6**), supporting the concentration-dependent trend observed in the static model (**Figure 3b**).

No significant changes in Brnoteplase thrombolytic efficacy were detected even after 3 months of storage at +4 °C (p=0.40), as opposed to alteplase (p=0.02), concluding the high shelf-life of Brnoteplase and its suitability for long-term storage applications (**Figure S12**, **Table S6**).

Fibrin gel penetration experiments (**Figure 3d**, **Table S7**) confirmed detectable interaction with fibrin for both alteplase and Brnoteplase, as both exhibited significant changes in penetration rates (p<0.001 at all distances) compared to the non-interacting control bovine serum albumin (BSA). Comparison of alteplase and Brnoteplase showed similar penetration rates for both proteins up to the distance of 0.50 mm (p=0.07 at 0.00 mm; p=0.08 at 0.25 mm; p=0.06 at 0.50 mm). However, a significantly higher amount of Brnoteplase penetrated beyond 0.75 mm (p=0.008 at 0.75 mm; p<0.001 at 1.00 mm), confirming the superior clot penetration potential identified during biochemical characterization (**Figure 2f**).

Altogether, *in vitro* analysis confirmed the strong potential of Brnoteplase due to multiple additional advantageous properties over alteplase: (i) thrombolytic efficacy increase with concentration, (ii) preserved stability for over 3 months at +4 °C, and (iii) faster penetration rate and greater distribution within the clot. Combined with its enhanced fibrin selectivity (**Figure 2c**) and PAI-1 inhibition resistance (**Figure 2e**), the Brnoteplase profile was indicating improved safety, recanalization potential, and biological half-life, which would enable bolus administration and increased dosing in the subsequent *in vivo* testing.

### Animal testing verifies increased Brnoteplase efficiency and safety *in vivo*

Our selected variant Brnoteplase and approved drugs alteplase and tenecteplase were tested *in vivo* for thrombolysis effectivity and safety in rat models (**Figure 4**). Thrombolytic effectivity was assessed by the lysis rate of administered thrombi and the area under the curve (AUC) of lytic curves. The Brnoteplase group (2.5 mg/kg, bolus) showed a lysis rate of 1.3 ± 0.4 %/min (**Figure 4a**, **Table S8**), which was the highest among groups when compared to 0.9 ± 0.5 %/min lysis rate for the alteplase (0.9 mg/kg, continuous rate infusion) and 1.1 ± 0.5 %/min for the tenecteplase (0.25 mg/kg, bolus) groups. Similarly, the AUC in Brnoteplase-treated rats was 2350 ± 720, exceeding the values observed in the alteplase (1520 ± 810) and tenecteplase (2180 ± 1060) groups (**Figure 4a**, **Table S8**). Statistical analysis confirmed that Brnoteplase demonstrated superior thrombolytic efficacy at all tested doses (0.9, 2.5, and 5.0 mg/kg; **Figure S13**, **Table S8**) by providing significantly higher lysis rates and AUC values than both control and alteplase (0.9 mg/kg, continuous rate infusion) groups (p ≤ 0.01). Interestingly, Brnoteplase (0.9 mg/kg) demonstrated activity at the level of untreated control when applied in the continuous rate infusion mode (**Table S8**).

**Figure 4:**
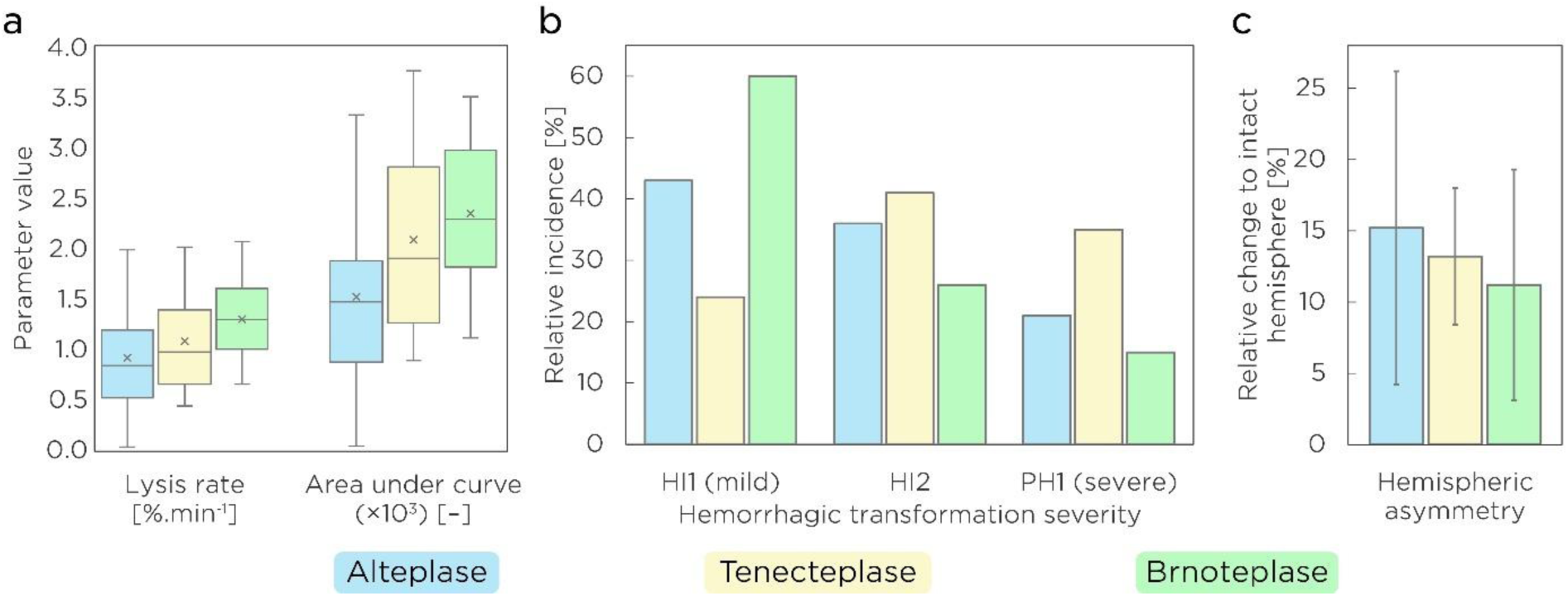
*In vivo* analysis of alteplase (0.9 mg/kg, continuous rate infusion), tenecteplase (0.25 mg/kg, bolus), and Brnoteplase (2.5 mg/kg, bolus) performance in animal rat models. (**a**) Comparison of the thrombolytic effectivity of alteplase (n=16), tenecteplase (n=12), and Brnoteplase (n=12), expressed as lysis rate (% of the clot dissolved per minute) and the area under the curve (AUC) of the lytic curve. Box plots illustrate mean values (cross), median (line), interquartile range (box), and minimum/maximum values (whiskers). (**b**) Proportion of hemorrhagic transformation severities according to ECASS II scoring system^42^ after administering alteplase (n=22), tenecteplase (n=24), or Brnoteplase (n=22). For a detailed description, see the Methods section. (**c**) Comparison of hemispheric asymmetry of rat brains expressed as a relative change to an intact hemisphere after administering alteplase (n=22), tenecteplase (n=24), or Brnoteplase (n=22). Bar plots illustrate mean values (bars) and standard errors (whiskers).

The safety study assessed the overall incidence of hemorrhagic transformation (HT) and its severity according to ECASS II criteria^42^ to evaluate the safety profile of each thrombolytic agent (**Figure 4b**, **Figure S14**, **Table S9**). Given the experimental model, in which the blood-brain barrier was damaged, and collaterals collapsed due to prolonged ischemia, bleeding could not occur without recanalization^48^, making overall HT incidence an indicator of successful recanalization. HT occurred most frequently (86 %) in the Brnoteplase group (2.5 mg/kg), followed by the 71 % incidence in the tenecteplase (0.25 mg/kg), and 64 % in the alteplase (0.9 mg/kg) groups, being positively correlated with recanalization frequencies (87 %, 68 %, and 56 %, respectively; **Figure S14**, **Table S9**). Among HT subtypes, Brnoteplase treatment led predominantly to mild petechial hemorrhages (HI1), occurring in 60 % of treated rats, while only 15 % resulted in severe hemorrhages (PH1). These results substantially differed from both alteplase (43 % HI1 vs. 21 % PH1) and tenecteplase (24 % HI1 vs. 35 % PH1) groups where the proportion of severe PH1 was more prevalent (**Figure 4b**, **Table S9**).

Hemispheric asymmetry analysis (**Figure 4c**, **Table S9**) yielded the lowest value of 11 ± 8 % for Brnoteplase-treated rats (2.5 mg/kg), followed by 13 ± 5 % for tenecteplase-treated (0.25 mg/kg) and 15 ± 11 % for alteplase-treated (0.9 mg/kg) groups. The lowest asymmetry in the Brnoteplase group is coupled with the highest overall HT incidence, reflecting the highest recanalization frequency (**Figure S14**, **Table S9**), and the predominance of the least dangerous HT grade (HI1). Together with the lowest observed proportion of severe HT (PH1; **Figure 4b**), this outcome indicates that Brnoteplase treatment was associated with reduced brain swelling and infarct expansion compared to the administration of alteplase or tenecteplase, suggesting a potentially more favorable safety profile.

### Rat plasma analysis corroborates superior Brnoteplase properties

Rat plasma collected at the end of the thrombolysis effectivity experiments was analyzed to determine circulating levels of plasminogen, plasmin, and the administered thrombolytic (**Figure 5**, **Table S10**). Residual thrombolytic concentrations provided information about residence time and half-life (**Figure 5a**, **Table S10**). The results yielded 37-, 11-, and 30-fold increased levels of active Brnoteplase over alteplase at doses of 0.9, 2.5, and 5.0 mg/kg, respectively. Notably, the residual active Brnoteplase concentration after administering the 0.9 mg/kg dose (0.22 ± 0.06 mg/L) was significantly higher even than the alteplase concentration after administering 5.0 mg/kg (0.09 ± 0.01 mg/L). The distinct administration regimes—bolus for Brnoteplase vs. continuous rate infusion for alteplase—underscore this difference and support that the improved inhibition resistance of Brnoteplase (**Figure 2e**) and potentially alleviated binding to clearance receptors (**Table 3**) substantially extended its biological half-life.

**Figure 5:**
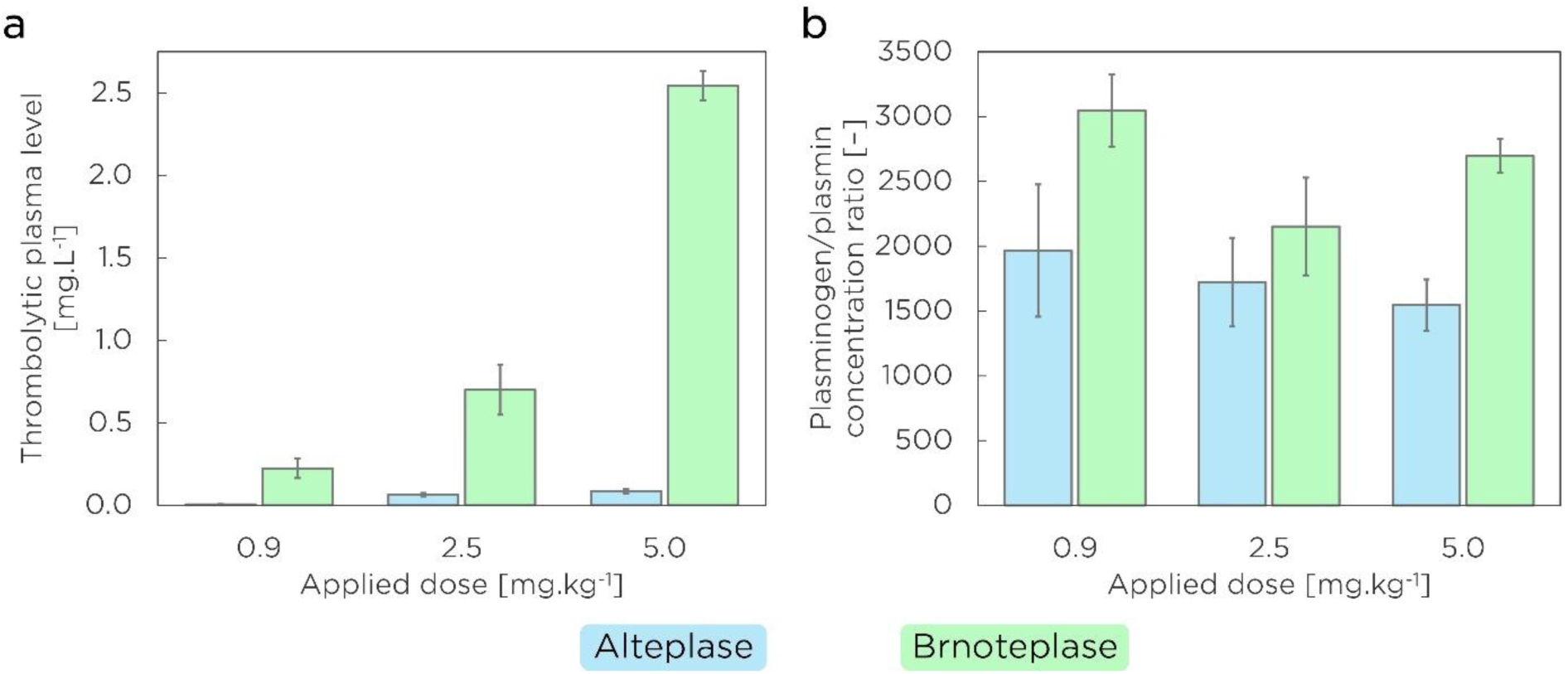
Determination of pharmacological marker levels in rat plasma samples collected at the end of the thrombolysis effectivity experiments. The bars represent mean values and the whiskers correspond to standard errors. The data were collected for alteplase applied as continuous rate infusion (n=7, 6, and 6 for the doses 0.9, 2.5, and 5.0 mg/kg, respectively) and Brnoteplase applied as bolus (n=16, 13, and 5 for the doses 0.9, 2.5, and 5.0 mg/kg, respectively). (**a**) Comparison of residual active concentrations of administered thrombolytics as an indicator of the protein residence time and biological half-life. (**b**) Comparison of residual plasminogen-to-plasmin ratios as an indicator of thrombolytic selectivity and safety potential.

Plasminogen and plasmin levels were quantified to evaluate the thrombolytic pharmacological profile. Substantial systemic plasminemia is associated with increased hemorrhagic risk, whereas preserving high plasminogen levels and restricting plasmin generation to occluded areas promises more favorable treatment outcomes^17^. Accordingly, the plasminogen/plasmin ratio serves as an indicator of thrombolytic selectivity and safety potential. Across all tested doses (0.9, 2.5, and 5.0 mg/kg), Brnoteplase-treated rats exhibited a consistent trend of higher plasminogen/plasmin ratios than the alteplase group (**Figure 5b**, **Table S10**). Furthermore, Brnoteplase treatment at the dose of 5.0 mg/kg retained a comparable or even higher ratio (2700 ± 130) relative to the alteplase group at the lower, clinically relevant dose of 0.9 mg/kg (1970 ± 510). Together with the significantly increased *in vivo* thrombolytic efficacy of Brnoteplase (**Figure 4a**), these findings corroborate previously observed enhanced selectivity (**Figure 2c**) and favorable safety profile (**Figure 4bc**) of Brnoteplase.

## Discussion

In this study, we established a multi-objective enzyme engineering pipeline that integrates *in silico* rational design, ancestral sequence reconstruction, database-guided selection, with biochemical characterization, *in vitro* clot lysis and penetration assays, and *in vivo* animal models. This workflow enables generation and systematic prioritization of thrombolytic candidates, focusing on multiple therapeutic parameters simultaneously. Using this approach, we identified Brnoteplase, a novel thrombolytic with improved pharmacological properties, and it represents a promising candidate for future preclinical and clinical evaluation.

Biochemical characterization revealed that Brnoteplase exhibited (i) markedly enhanced fibrin selectivity (80-fold higher than alteplase), (ii) superior resistance to PAI-1 inhibition (4-fold higher), and (iii) increased clot penetration potential (4-fold higher). These parameters are critical determinants of safer and more effective thrombolysis and provide a mechanistic basis for the favorable outcomes observed in subsequent *in vitro* and *in vivo* experiments.

The pronounced fibrin selectivity of Brnoteplase ensures activation of plasminogen predominantly within occluded regions with fibrin-containing thrombi, while minimizing systemic plasminemia associated with hemorrhagic risk. As a result, Brnoteplase maintains a higher plasminogen/plasmin ratio relative to alteplase *in vivo*, as confirmed by ELISA analysis of rat plasma collected after thrombolysis. Additionally, the *in silico* design of Brnoteplase aimed to reduce affinity for LRP1 and NMDAR which is expected to mitigate off-target central nervous system interactions and thereby further enhance the safety profile^15,16^. All these findings were corroborated *in vivo*, where the Brnoteplase-treated group exhibited a lower proportion of severe hemorrhagic transformations compared to alteplase and tenecteplase.

Enhanced inhibition resistance, another major characteristic of Brnoteplase, aligns with expectations from the rational design phase. This variant incorporates mutations previously shown to reduce interaction with PAI-1, a primary inhibitor of tissue plasminogen activator in human blood. Together with mutations designed to minimize binding to clearance receptors such as LRP1 and mannose receptors, these combined features were intended to modulate protein pharmacokinetics^6,49,50^, and this expectation was confirmed *in vivo*. Rat plasma samples collected at the end of thrombolysis experiments revealed significantly higher residual levels of active Brnoteplase compared to alteplase, indicating an extended biological half-life. This enabled efficient bolus administration, a clinically meaningful advantage over alteplase.

Finally, superior clot penetration potential of Brnoteplase in biochemical assays was derived from decreased fibrin affinity. This assumption was validated by *in vitro* assays directly monitoring thrombolytics motion through fibrin matrix gels. Compared to alteplase, Brnoteplase exhibited a faster penetration rate and greater distribution within the clot. This facilitates lysis from within the clot rather than relying solely on surface degradation, which is characteristic of conventional thrombolytics. By promoting more uniform and deeper fibrinolysis, Brnoteplase may accelerate recanalization, particularly in large or densely structured thrombi.

In addition to these mechanistic advantages, Brnoteplase demonstrated a favorable dose–response profile *in vitro*. Thrombolytic efficacy progressively increased with concentration until plateau, whereas alteplase exhibited an early peak with a decline at elevated doses. Combined with the Brnoteplase selectivity and reduced affinity for receptors, this profile suggests that higher dosing could enhance thrombolytic potency without compromising safety and hemorrhagic complications. Consistent with these findings, *in vivo* experiments in rat models confirmed that Brnoteplase achieved significantly higher thrombolysis rates and recanalization frequency across all tested doses compared to alteplase. Importantly, safety assessments demonstrated the lowest incidence of severe hemorrhagic transformation and minimal hemispheric asymmetry in Brnoteplase-treated animals relative to both alteplase and tenecteplase. Nevertheless, given the low overall occurrence of hemorrhagic events in the tested cohorts, larger-scale studies will be required to confirm statistical significance of the safety study outcome.

Considering that Brnoteplase incorporates several mutations originally introduced in tenecteplase, it is not unexpected that they share a similar profile in biochemical testing, namely fibrin selectivity and PAI-1 resistance. However, Brnoteplase also demonstrated significantly greater fibrin clot penetration and a lower rate of severe hemorrhagic transformation, positioning it as a potentially advantageous alternative within the current state-of-the-art thrombolytics.

In summary, we establish an enzyme engineering workflow that integrates multi-objective design, prioritization, and multilevel functional validation for the development of improved thrombolytic biocatalysts. Brnoteplase exemplifies the outcome of this approach, combining enhanced fibrin selectivity, increased resistance to inhibition, prolonged functional lifetime, and improved substrate accessibility within fibrin matrices. These properties collectively enhance effective catalytic performance under physiologically relevant conditions, resulting in improved thrombolytic activity and a favorable safety profile in preclinical *in vivo* models. Beyond the specific case of Brnoteplase, our work provides a generalizable framework for optimizing proteolytic enzymes operating in complex biological environments. The proposed approach builds upon existing enzyme catalysis methods and emphasizes the potential of combining rational and evolutionary design to create next-generation therapeutics biocatalysts., where effective catalysis requires coordinated optimization of multiple parameters. While further validation in advanced models is required, the presented approach expands current strategies in enzyme catalysis and highlights the potential of integrating rational and evolutionary design to develop next-generation therapeutic biocatalysts.

## Author Contribution Statement

**J.M and D.B.** performed the *in silico* analysis and selection of the initial thrombolytic pool. **Pe.Ka. and T.B.** produced the designed thrombolytic enzymes. **M.T. and V.S.** performed the biochemical characterization of selected proteins and ELISA measurements of rat plasma samples and analyzed the corresponding data. **S.T., M.P., Pa.Ki., and J.V.** performed the *in vitro* thrombolysis and fibrin penetration assays and analyzed the corresponding data. **P.S., J.H., E.B., A.D.A., J.B., M.K., and J.O.** performed the *in vivo* animal testing and the corresponding data analysis and statistical evaluation. **M.T., V.S.** drafted the initial version of the manuscript with the help of **J.M., S.T., and J. V.**. **J.D., Z.P., D.B., M.M., J.V., L.K., and R.M.** conceived and designed the study, supervised the project, and acquired funding. **All authors** reviewed and interpreted the data, critically revised the manuscript, and approved the final version.

## Acknowledgements

We thank the laboratory technicians Marketa Stuchla, Veronika Novakova, Irena Halikova, and Klara Markova for their assistance with the preparation of common reagents and sample storage.

## References

1. Feigin VL, Brainin M, Norrving B, et al. World Stroke Organization: Global stroke fact sheet 2025. Int J Stroke. 2025;20(2):132–144. doi:10.1177/17474930241308142

2. Aguiar de Sousa D, von Martial R, Abilleira S, et al. Access to and delivery of acute ischaemic stroke treatments: A survey of national scientific societies and stroke experts in 44 European countries. Eur Stroke J. 2019;4(1):13–28. doi:10.1177/2396987318786023

3. Xiong Y, Wakhloo AK, Fisher M. Advances in acute ischemic stroke therapy. Circ Res. 2022;130(8):1230–1251. doi:10.1161/CIRCRESAHA.121.319948

4. Yaghi S, Willey JZ, Cucchiara B, et al. Treatment and outcome of hemorrhagic transformation after intravenous alteplase in acute ischemic stroke: A scientific statement for healthcare professionals from the American Heart Association/American Stroke Association. Stroke. 2017;48(12):e343–e361. doi:10.1161/STR.0000000000000152

5. Mosconi MG, Paciaroni M. Treatments in ischemic stroke: Current and future. Eur Neurol. 2022;85(5):349–366. doi:10.1159/000525822

6. Mican J, Toul M, Bednar D, Damborsky J. Structural biology and protein engineering of thrombolytics. Comput Struct Biotechnol J. 2019;17:917–938. doi:10.1016/j.csbj.2019.06.023

7. Potla N, Ganti L. Tenecteplase vs alteplase for acute ischemic stroke: A systematic review. Int J Emerg Med. 2022;15(1):1. doi:10.1186/s12245-021-00399-w

8. Burwell JM, Howay JR, Wasko L, et al. Tenecteplase is here: Navigating the shift of a stroke thrombolytic in the United States prior to FDA approval: A mini-review on rationale, barriers, and pathways. Front Neurol. 2025;16:1563423. doi:10.3389/fneur.2025.1563423

9. Rodriguez M, Sidebottom C, Wells DA, et al. A systematic review of the efficacy and safety of tenecteplase versus alteplase in acute ischemic stroke: A time to pass the torch. Stroke Vasc Interv Neurol. 2024;4(4):e001110. doi:10.1161/SVIN.123.001110

10. Ma Y, Xiang H, Busse JW, et al. Tenecteplase versus alteplase for acute ischemic stroke: A systematic review and meta-analysis of randomized and non-randomized studies. J Neurol. 2024;271(5):2309–2323. doi:10.1007/s00415-024-12243-1

11. Rose D, Cavalier A, Kam W, et al. Complications of intravenous tenecteplase versus alteplase for the treatment of acute ischemic stroke: A systematic review and meta-analysis. Stroke. 2023;54(5):1192–1204. doi:10.1161/STROKEAHA.122.042335

12. Warach SJ, Dula AN, Milling TJ. Tenecteplase thrombolysis for acute ischemic stroke. Stroke. 2020;51(11):3440–3451. doi:10.1161/STROKEAHA.120.029749

13. Meng X, Li S, Dai H, et al. Tenecteplase vs alteplase for patients with acute ischemic stroke: The ORIGINAL randomized clinical trial. JAMA. 2024;332(17):1437–1445. doi:10.1001/jama.2024.14721

14. Van de Werf FJ. The ideal fibrinolytic: Can drug design improve clinical results? Eur Heart J. 1999;20(20):1452–1458. doi:10.1053/euhj.1999.1659

15. Maier W, Bednorz M, Meister S, et al. LRP1 is critical for the surface distribution and internalization of the NR2B NMDA receptor subtype. Mol Neurodegener. 2013;8(1):25. doi:10.1186/1750-1326-8-25

16. Baron A, Montagne A, Cassé F, et al. NR2D-containing NMDA receptors mediate tissue plasminogen activator-promoted neuronal excitotoxicity. Cell Death Differ. 2010;17(5):860–871. doi:10.1038/cdd.2009.172

17. Nikitin D, Choi S, Mican J, et al. Development and testing of thrombolytics in stroke. J Stroke. 2021;23(1):12–36. doi:10.5853/jos.2020.03349

18. Sumbalova L, Stourac J, Martinek T, Bednar D, Damborsky J. HotSpot Wizard 3.0: Web server for automated design of mutations and smart libraries based on sequence input information. Nucleic Acids Res. 2018;46(W1):W356–W362. doi:10.1093/nar/gky417

19. Alford RF, Leaver-Fay A, Jeliazkov JR, et al. The Rosetta all-atom energy function for macromolecular modeling and design. J Chem Theory Comput. 2017;13(6):3031–3048. doi:10.1021/acs.jctc.7b00125

20. Prasad JM, Young PA, Strickland DK. High-affinity binding of the receptor-associated protein D1D2 domains with the low-density lipoprotein receptor-related protein (LRP1) involves bivalent complex formation: Critical roles of lysines 60 and 191. J Biol Chem. 2016;291(35):18430–18439. doi:10.1074/jbc.M116.744904

21. Freeman R, Niego B, R. Croucher D, Pedersen LO, Medcalf RL. t-PA, but not desmoteplase, induces plasmin-dependent opening of a blood-brain barrier model under normoxic and ischaemic conditions. Brain Res. 2014;1565:63–73. doi:10.1016/j.brainres.2014.03.027

22. Musil M, Khan RT, Beier A, et al. FireProt^ASR^: A web server for fully automated ancestral sequence reconstruction. Brief Bioinform. 2021;22(4):bbaa337. doi:10.1093/bib/bbaa337

23. Hon J, Borko S, Stourac J, et al. EnzymeMiner: Automated mining of soluble enzymes with diverse structures, catalytic properties and stabilities. Nucleic Acids Res. 2020;48(W1):W104–W109. doi:10.1093/nar/gkaa372

24. Toul M, Strunga A, Damborsky J, Prokop Z. Thrombolytic proteins profiling: High-throughput activity, selectivity, and resistance assays. FEBS Open Bio. doi:10.1002/2211-5463.70132

25. Toschi L, Bringmann P, Petri T, Donner P, Schleuning WD. Fibrin selectivity of the isolated protease domains of tissue-type and vampire bat salivary gland plasminogen activators. Eur J Biochem. 1998;252(1):108–112. doi:10.1046/j.1432-1327.1998.2520108.x

26. Harpaz D, Chen X, Francis CW, Marder VJ, Meltzer RS. Ultrasound enhancement of thrombolysis and reperfusion *in vitro*. J Am Coll Cardiol. 1993;21(6):1507–1511. doi:10.1016/0735-1097(93)90331-T

27. Prasad S, Kashyap RS, Deopujari JY, Purohit HJ, Taori GM, Daginawala HF. Development of an *in vitro* model to study clot lysis activity of thrombolytic drugs. Thromb J. 2006;4:14. doi:10.1186/1477-9560-4-14

28. Thalerová S, Pešková M, Kittová P, et al. Effect of apixaban pretreatment on alteplase-induced thrombolysis: An *in vitro* study. Front Pharmacol. 2021;12:740930. doi:10.3389/fphar.2021.740930

29. Víteček J, Wünschová AV, Thalerová S, et al. Factors influencing the efficacy of recombinant tissue plasminogen activator: Implications for ischemic stroke treatment. PLOS ONE. 2024;19(6):e0302269. doi:10.1371/journal.pone.0302269

30. Thalerová S, Wünschová AV, Kittová P, et al. A collateral circulation in ischemic stroke accelerates recanalization due to lower clot compaction. PLOS ONE. 2024;19(11):e0314079. doi:10.1371/journal.pone.0314079

31. Nikitin D, Mican J, Toul M, et al. Computer-aided engineering of staphylokinase toward enhanced affinity and selectivity for plasmin. Comput Struct Biotechnol J. 2022;20:1366–1377. doi:10.1016/j.csbj.2022.03.004

32. Diamond SL. Engineering design of optimal strategies for blood clot dissolution. Annu Rev Biomed Eng. 1999;1:427–461. doi:10.1146/annurev.bioeng.1.1.427

33. Acheampong P, and Ford GA. Pharmacokinetics of alteplase in the treatment of ischaemic stroke. Expert Opin Drug Metab Toxicol. 2012;8(2):271–281. doi:10.1517/17425255.2012.652615

34. Elnager A, Abdullah WZ, Hassan R, et al. *In vitro* whole blood clot lysis for fibrinolytic activity study using D-dimer and confocal microscopy. Adv Hematol. 2014;2014:814684. doi:10.1155/2014/814684

35. Rottenberger Z, Komorowicz E, Szabó L, et al. Lytic and mechanical stability of clots composed of fibrin and blood vessel wall components. J Thromb Haemost. 2013;11(3):529–538. doi:10.1111/jth.12112

36. Marcos-Contreras OA, Ganguly K, Yamamoto A, et al. Clot penetration and retention by plasminogen activators promote fibrinolysis. Biochem Pharmacol. 2013;85(2):216–222. doi:10.1016/j.bcp.2012.10.011

37. Longstaff C. Measuring fibrinolysis: From research to routine diagnostic assays. J Thromb Haemost. 2018;16(4):652–662. doi:10.1111/jth.13957

38. Longstaff C, Varjú I, Sótonyi P, et al. Mechanical stability and fibrinolytic resistance of clots containing fibrin, DNA, and histones. J Biol Chem. 2013;288(10):6946–6956. doi:10.1074/jbc.M112.404301

39. Morrow GB, Whyte CS, Mutch NJ. Functional plasminogen activator inhibitor 1 is retained on the activated platelet membrane following platelet activation. Haematologica. 2019;105(12):2824–2833. doi:10.3324/haematol.2019.230367

40. Vítečková Wünschová A, Novobilský A, Hložková J, et al. Thrombus imaging using 3D printed middle cerebral artery model and preclinical imaging techniques: Application to thrombus targeting and thrombolytic studies. Pharmaceutics. 2020;12(12):1207. doi:10.3390/pharmaceutics12121207

41. Kavkova M, Zikmund T, Kala A, et al. Contrast enhanced X-ray computed tomography imaging of amyloid plaques in Alzheimer disease rat model on lab based micro CT system. Sci Rep. 2021;11(1):5999. doi:10.1038/s41598-021-84579-x

42. Hacke W, Kaste M, Fieschi C, et al. Randomised double-blind placebo-controlled trial of thrombolytic therapy with intravenous alteplase in acute ischaemic stroke (ECASS II). The Lancet. 1998;352(9136):1245–1251. doi:10.1016/S0140-6736(98)08020-9

43. Bassel-Duby R, Jiang NY, Bittick T, et al. Tyrosine 67 in the epidermal growth factor-like domain of tissue-type plasminogen activator is important for clearance by a specific hepatic receptor. J Biol Chem. 1992;267(14):9668–9677.

44. López-Atalaya JP, Roussel BD, Ali C, et al. Recombinant *Desmodus rotundus* salivary plasminogen activator crosses the blood-brain barrier through a low-density lipoprotein receptor-related protein-dependent mechanism without exerting neurotoxic effects. Stroke. 2007;38(3):1036–1043. doi:10.1161/01.STR.0000258100.04923.84

45. Keyt BA, Paoni NF, Refino CJ, et al. A faster-acting and more potent form of tissue plasminogen activator. Proc Natl Acad Sci U S A. 1994;91(9):3670–3674. doi:10.1073/pnas.91.9.3670

46. Goulay R, Naveau M, Gaberel T, Vivien D, Parcq J. Optimized tPA: A non-neurotoxic fibrinolytic agent for the drainage of intracerebral hemorrhages. J Cereb Blood Flow Metab. 2018;38(7):1180–1189. doi:10.1177/0271678X17719180

47. Narita M, Bu G, Herz J, Schwartz AL. Two receptor systems are involved in the plasma clearance of tissue-type plasminogen activator (t-PA) *in vivo*. J Clin Invest. 1995;96(2):1164–1168. doi:10.1172/JCI118105

48. Ma J, Ma Y, Shuaib A, Winship IR. Improved collateral flow and reduced damage after remote ischemic perconditioning during distal middle cerebral artery occlusion in aged rats. Sci Rep. 2020;10(1):12392. doi:10.1038/s41598-020-69122-8

49. Biessen EAL, van Teijlingen M, Vietsch H, et al. Antagonists of the mannose receptor and the LDL receptor–related protein dramatically delay the clearance of tissue plasminogen activator. Circulation. 1997;95(1):46–52. doi:10.1161/01.CIR.95.1.46

50. Chandler WL, Alessi MC, Aillaud MF, Henderson P, Vague P, Juhan-Vague I. Clearance of tissue plasminogen activator (TPA) and TPA/plasminogen activator inhibitor type 1 (PAI-1) complex. Circulation. 1997;96(3):761–768. doi:10.1161/01.CIR.96.3.761

