## Supporting information for "Multi-Objective Engineering of Fibrin-Selective Thrombolytic Proteases with Enhanced Biocatalytic Efficiency and Inhibition Resistance"

* Corresponding authors

### Supplementary figures

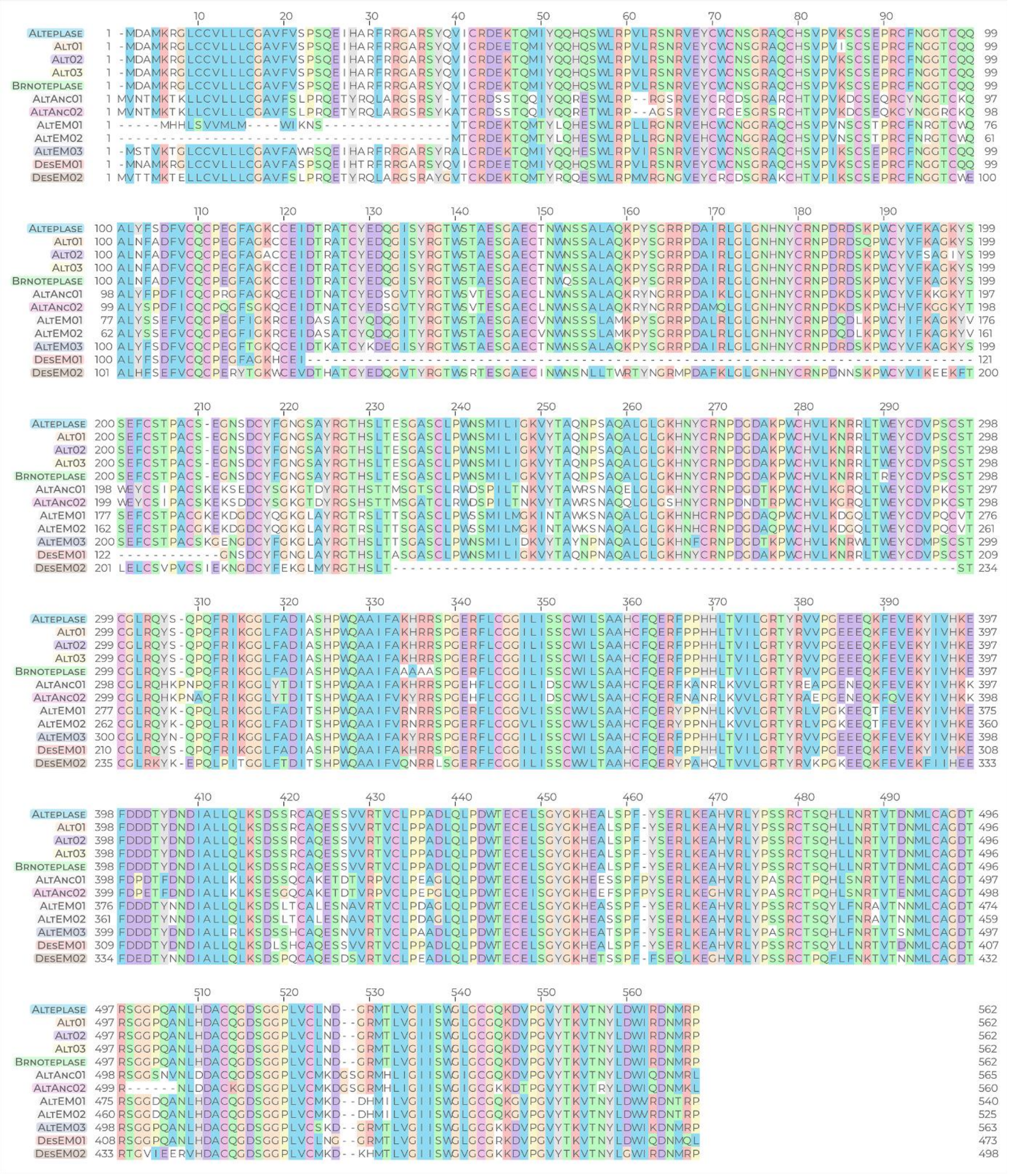

**Figure S1: Sequence alignment of protein sequences of thrombolytics selected during the *in silico* design step.** The selection includes 11 different proteins and the alignment is complemented with the reference alteplase sequence.

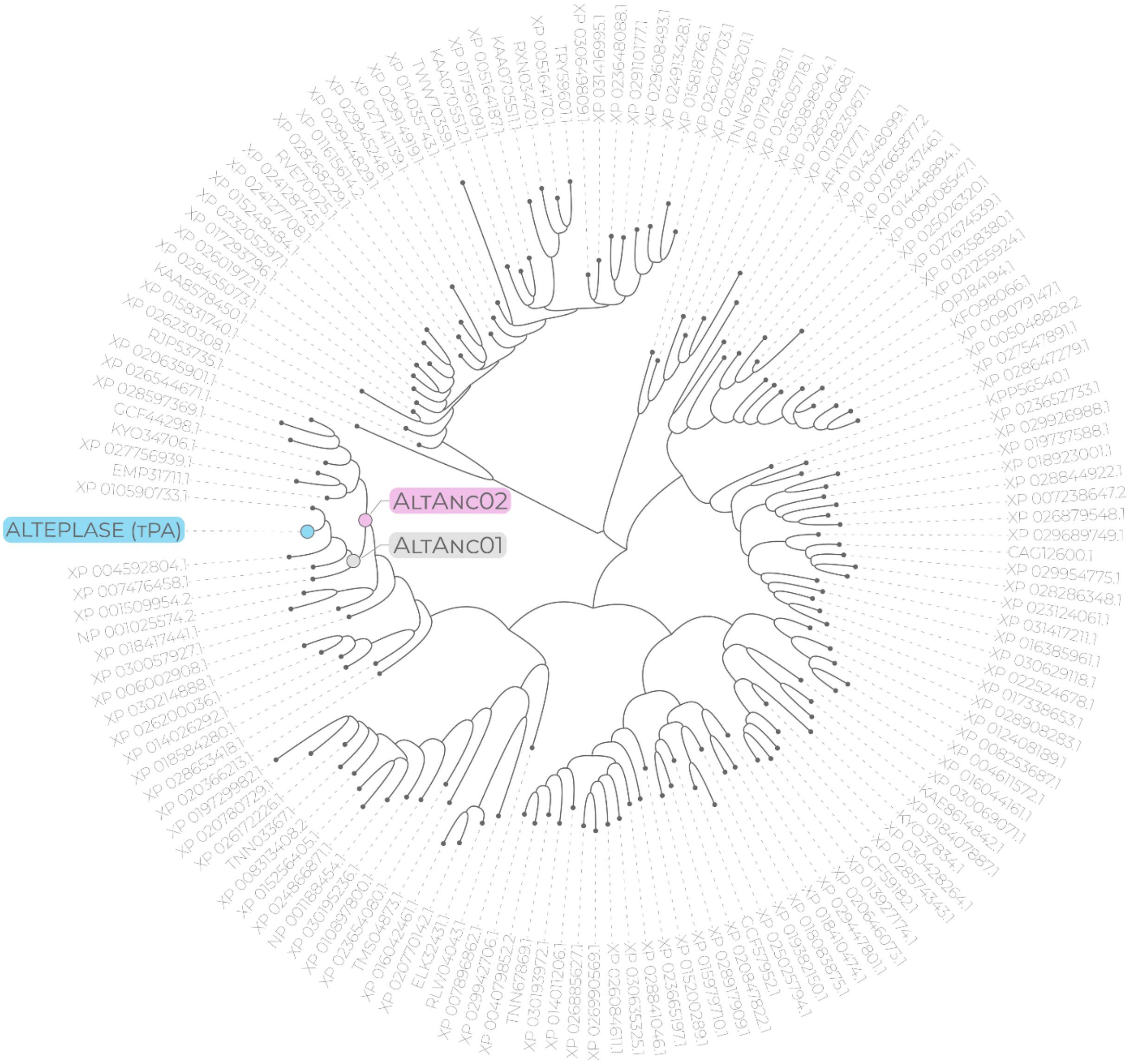

**Figure S2: Reconstructed phylogenetic tree of sequences related to alteplase (tPA).** The query sequence of alteplase (tPA) is highlighted in blue and the nodes corresponding to the two reconstructed ancestral sequences selected for biochemical characterization (AltAnc01 and AltAnc02) are shown in pink and grey, respectively. AltAnc01 is ancestral to 5 modern-day sequences (including alteplase) annotated as plasminogen activators while AltAnc02 is ancestral to 12 sequences (11 annotated as plasminogen activators + 1 hypothetical protein predicted to be a plasminogen activator). Codes of the other sequences correspond to their accession numbers in the corresponding databases.

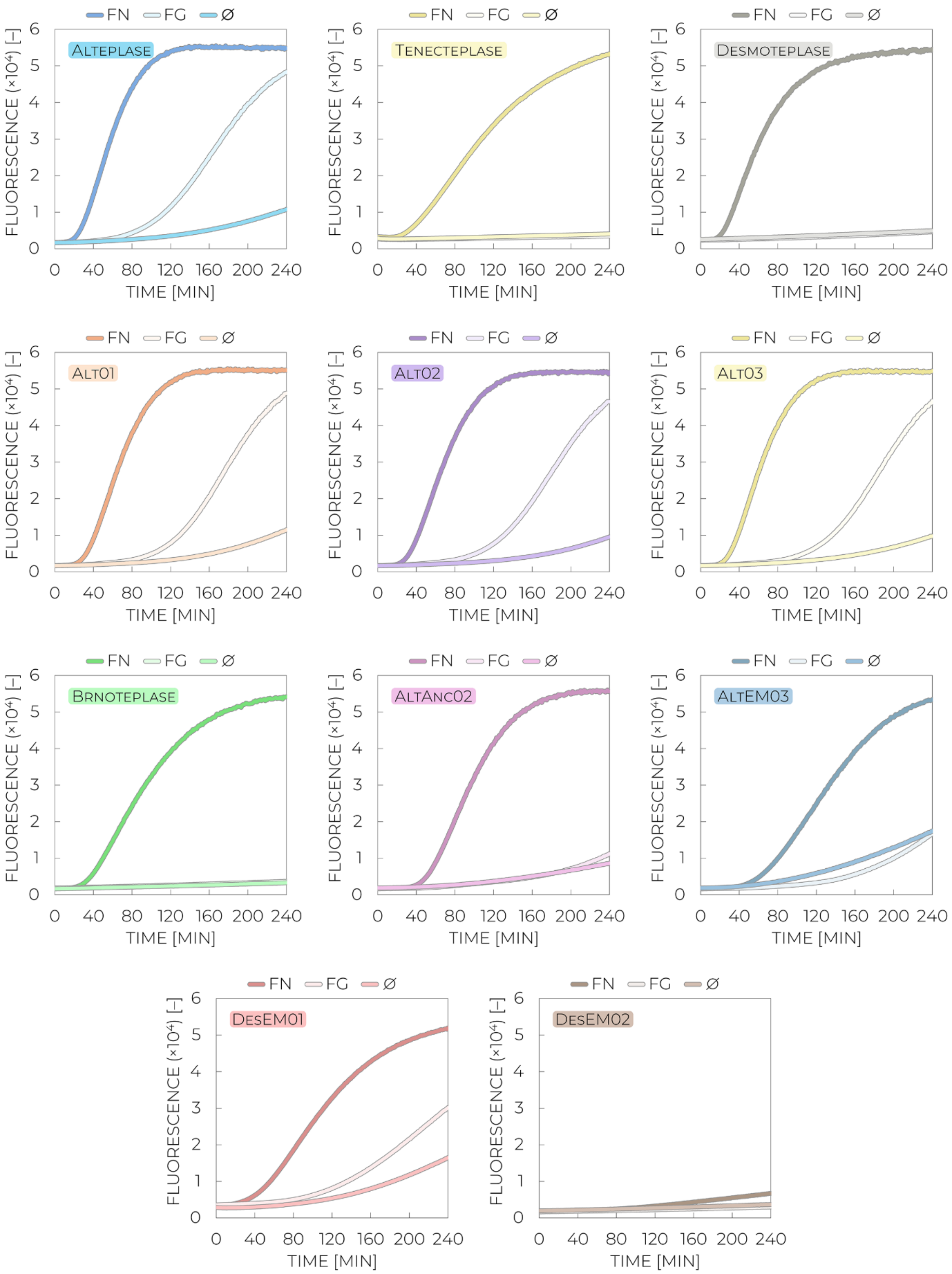

**Figure S3: Representative kinetic curves of plasminogen activation enzymatic activities with and without stimulation collected for variants tested in the biochemical characterization step.** The activities were measured in the absence of any stimulant (∅) and in the presence of fibrinogen (Fg) and fibrin (Fn). The experiments were performed at 37 °C in physiological PBS buffer pH 7.4 containing 1 mM $CaCl_2$, 0.0035 % L-arginine, and 0.01 % Tween 80.

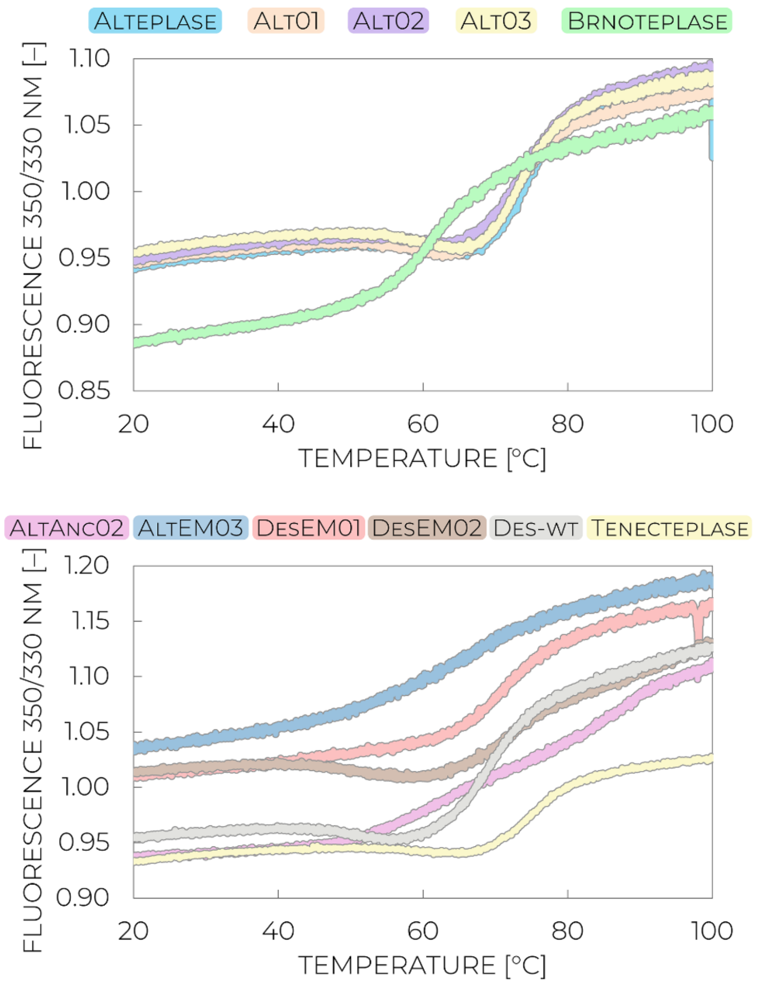

**Figure S4: Raw unfolding curves collected in the thermostability biochemical assay.** The curves show changes in native tryptophan fluorescence with increasing temperature as a result of protein unfolding and solvent exposure of tryptophan residues. The determined melting temperatures ($T_m$) correspond to inflection points of the sigmoid curves. The experiments were performed in physiological PBS buffer pH 7.4 containing 1 mM $CaCl_2$, 0.0035 % L-arginine, and 0.01 % Tween 80 by applying the temperature ramp of 20–100 °C.

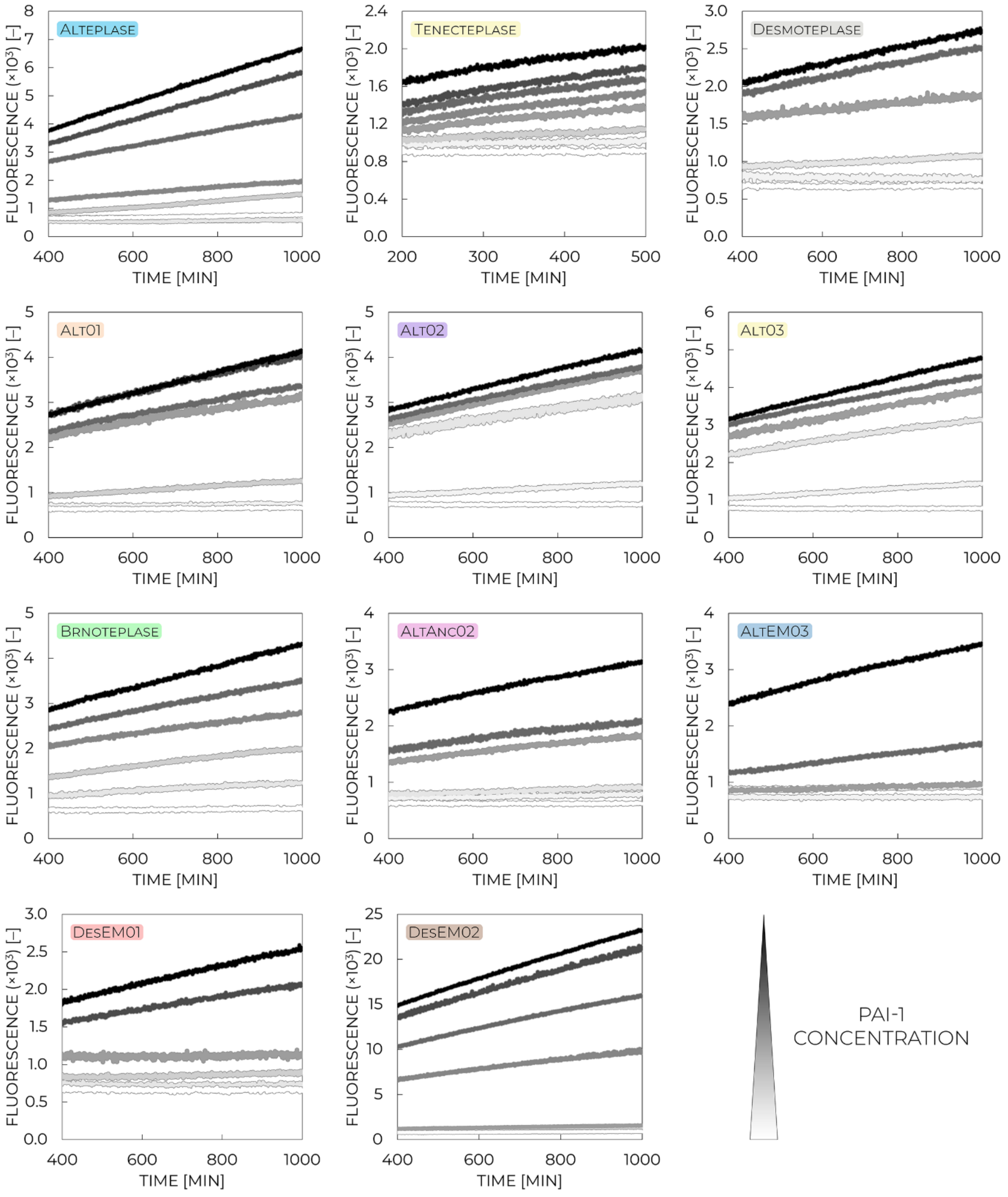

**Figure S5: Raw kinetic data collected in the PAI-1 inhibition resistance biochemical assay.** The curves correspond to residual amidolytic activities of the corresponding tested thrombolytic variants upon their incubation with varied plasminogen activator inhibitor-1 (PAI-1) concentrations, ranging from 0 to 1.86 µM. The residual activities are proportional to the amount of the non-inhibited/active thrombolytic fraction, allowing determination of inhibition constants based on concentration dependence analyses (**Figure S6**). The experiments were performed at 37 °C in physiological PBS buffer pH 7.4 containing 1 mM $CaCl_2$, 0.0035 % L-arginine, and 0.01 % Tween 80.

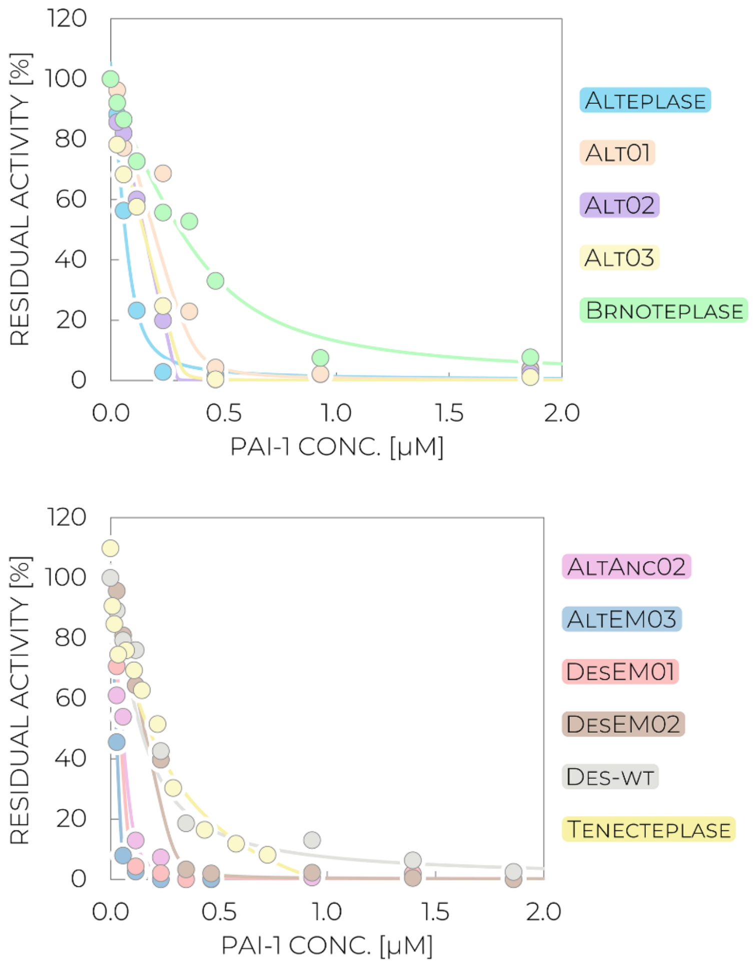

**Figure S6: PAI-1 concentration dependence analysis of residual activities of thrombolytic enzymes tested in the biochemical characterization step.** Fitting with the equation for a tightly bound inhibitor enables determination of the plasminogen activator inhibitor-1 (PAI-1) inhibition constants ($IC_{50}$), providing information about the concentration at which the activity drops to 50 %, i.e., information about the inhibition resistance of the tested thrombolytic variants. The experimental data are shown with circles while solid lines represent the best fit. The experimental data points were derived from experiments performed at 37 °C in physiological PBS buffer pH 7.4 containing 1 mM $CaCl_2$, 0.0035 % L-arginine, and 0.01 % Tween 80.

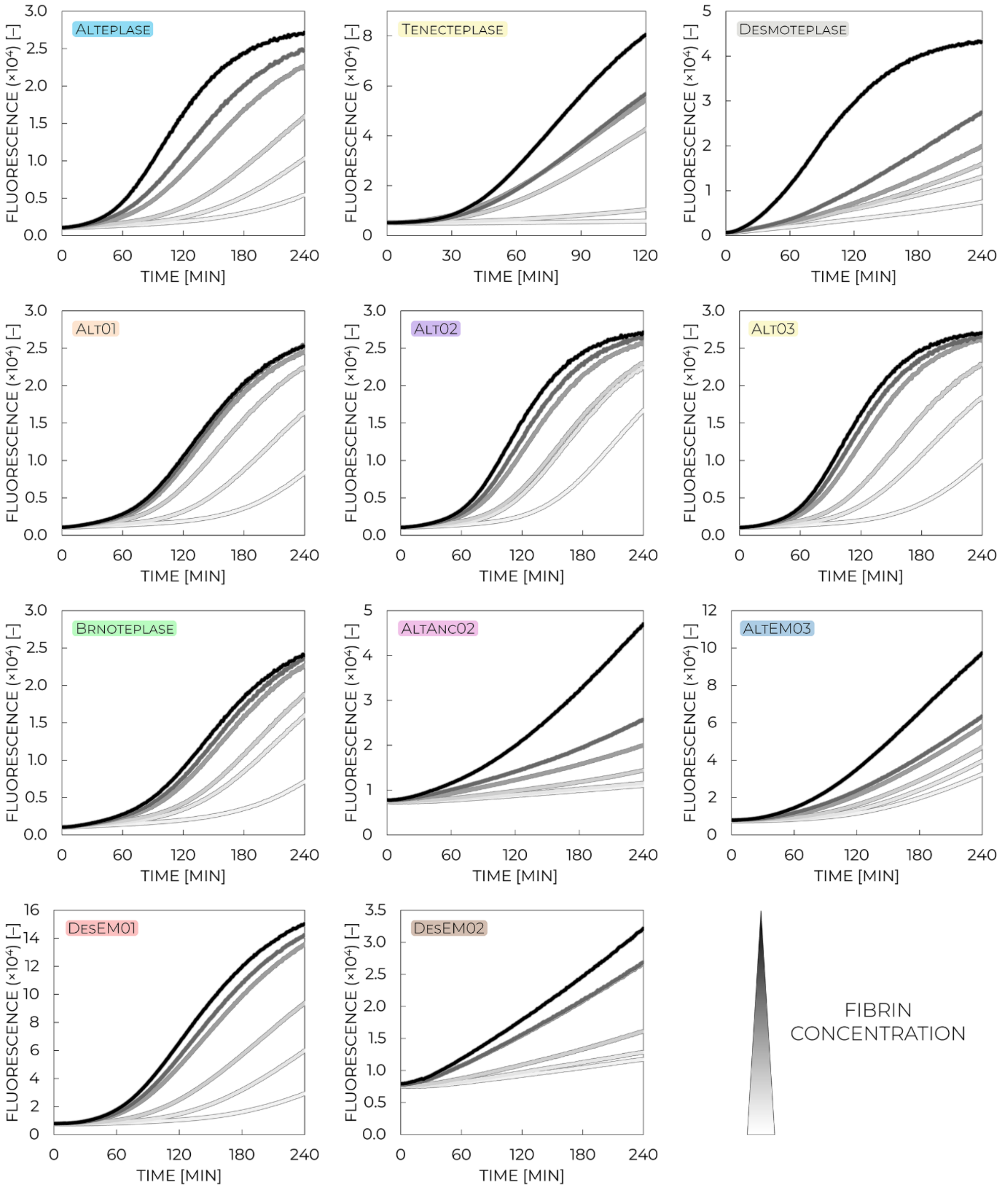

**Figure S7: Raw kinetic data collected in the fibrin binding biochemical assay.** The curves correspond to residual plasminogen activation activities of the corresponding tested thrombolytic variants upon their incubation with fibrin clots containing varied fibrin concentrations, ranging from 0 to 37.7 µM. The residual activities are proportional to the amount of the fibrin-unbound thrombolytic fraction, allowing determination of binding affinities based on concentration dependence analyses (**Figure S8**). The experiments were performed at 37 °C in physiological PBS buffer pH 7.4 containing 1 mM $CaCl_2$, 0.0035 % L-arginine, and 0.01 % Tween 80.

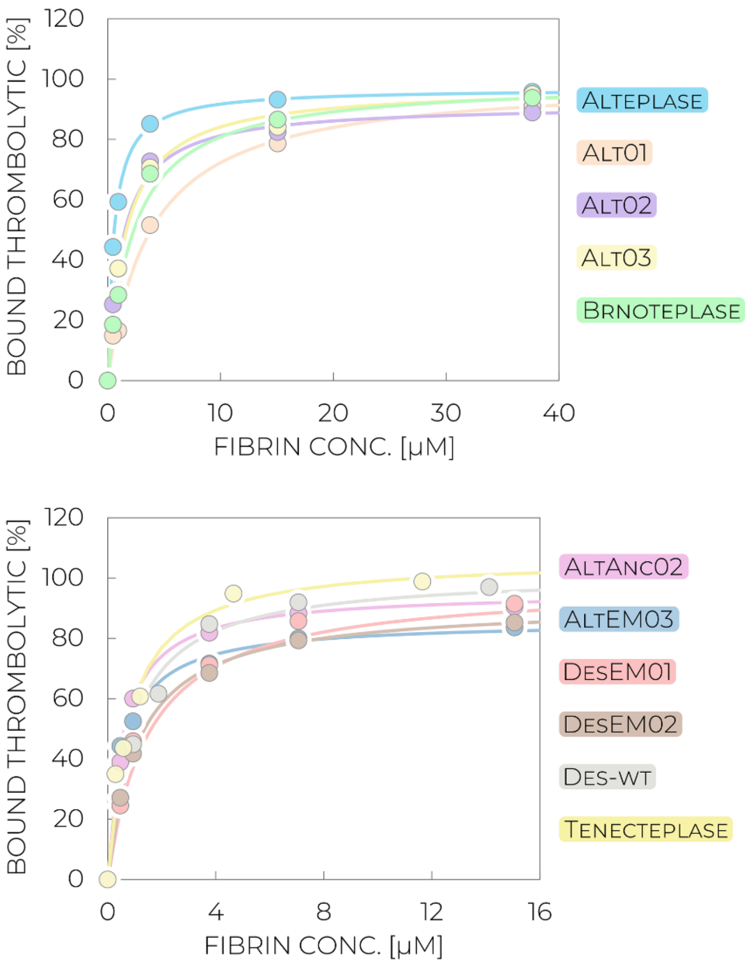

**Figure S8: Fibrin concentration dependence analysis of the amount of bound thrombolytic enzymes tested in the biochemical characterization step.** Fitting with the typical hyperbolic equation for an equilibrium thrombolytic binding to fibrin clots enables determination of the fibrin dissociation constants ($K_d$), providing information about the concentration at which half of the thrombolytic molecules are bound to the clot, i.e., information about the affinity and clot penetration potential of the tested thrombolytic variants. The experimental data are shown with circles while solid lines represent the best fit. The experimental data points were derived from experiments performed at 37 °C in physiological PBS buffer pH 7.4 containing 1 mM $CaCl_2$, 0.0035 % L-arginine, and 0.01 % Tween 80.

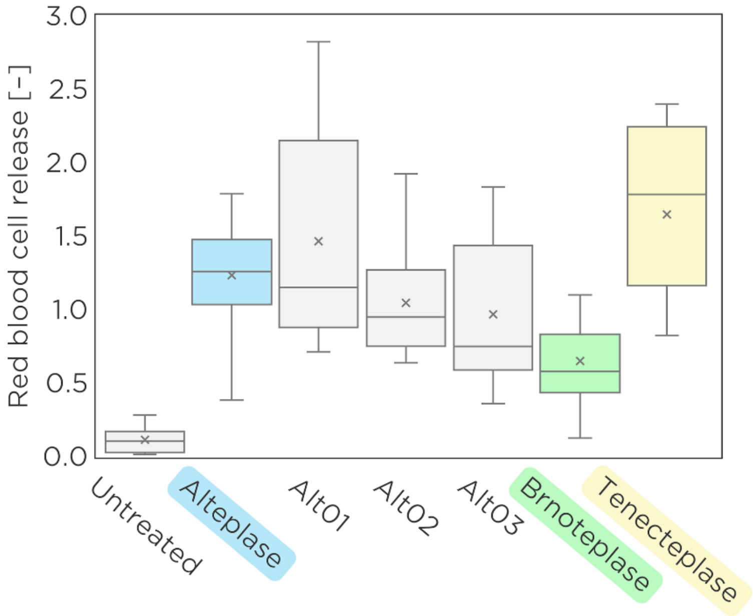

**Figure S9:** ***In vitro* analysis of clot lysis efficacy in the static model with semi-synthetic clots expressed as red blood cell release.** The comparison includes untreated group, reference thrombolytics alteplase (blue) and tenecteplase (yellow), and new tested variants Brnoteplase (green), Alt01, Alt02, and Alt03 (grey). Box plots illustrate mean values (cross), median (line), interquartile range (box), and minimum/maximum values (whiskers).

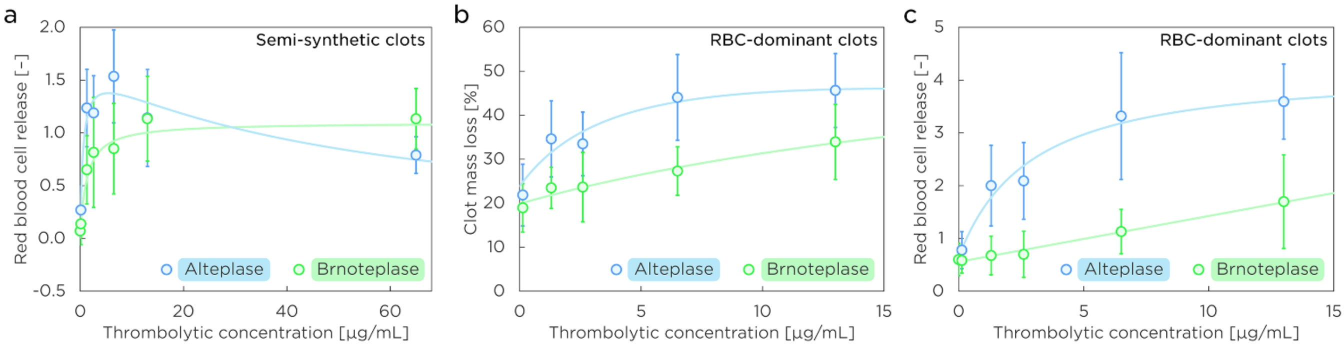

**Figure S10: Concentration dependence profiles of clot lysis efficacy in different regimes of the static model.** The dependence monitors efficacy trends with increasing applied concentration of a tested thrombolytic, i.e., alteplase (blue) or Brnoteplase (green). Scatter plots show mean values (circle) and standard errors (whiskers). (**a**) Concentration dependence obtained with semi-synthetic clots, expressed as red blood cell (RBC) release. (**b**) Concentration dependence obtained with RBC-dominant clots, expressed as clot mass loss. (**c**) Concentration dependence obtained with RBC-dominant clots, expressed as RBC release.

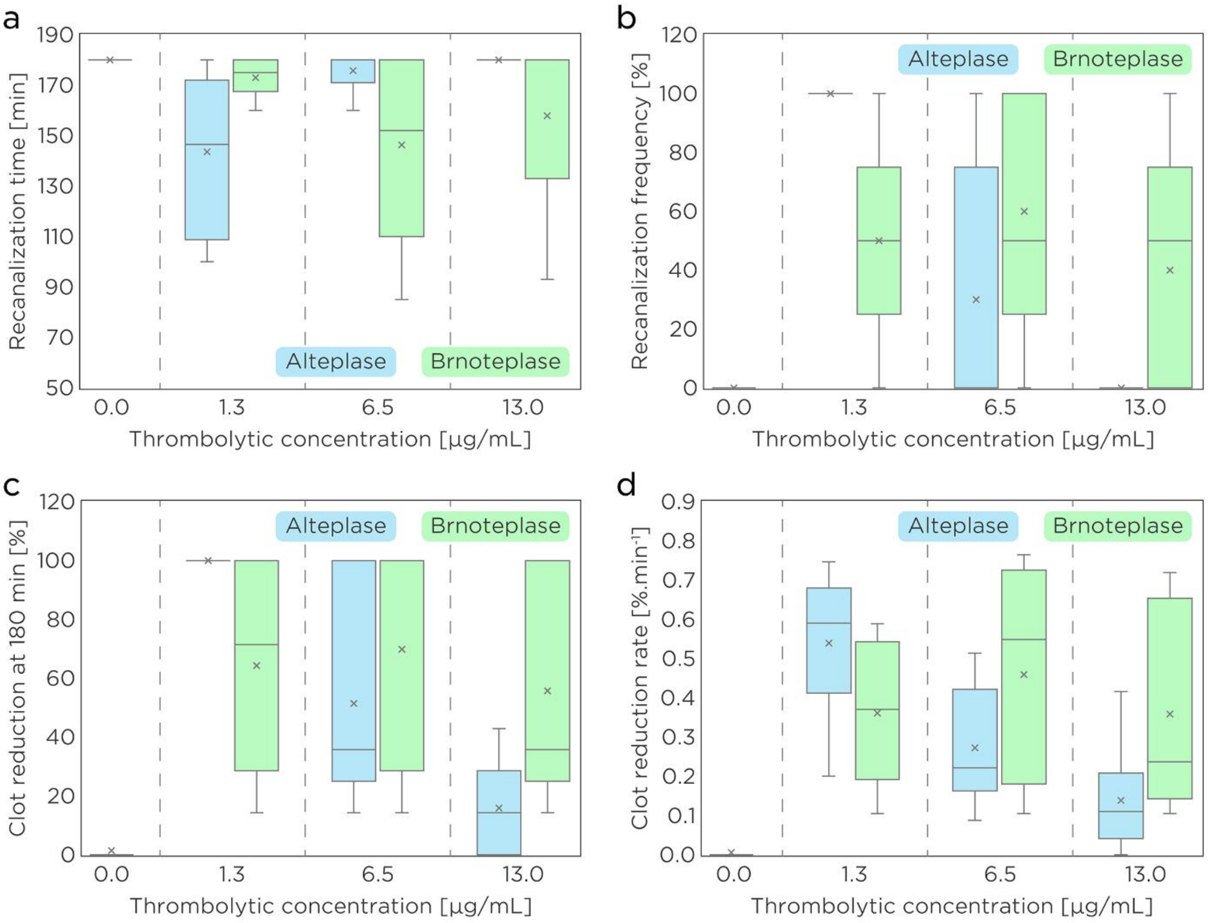

**Figure S11: *In vitro* analysis of clot lysis efficacy and concentration trends in the flow model with semi-synthetic clots.** The comparison includes untreated group (grey; concentration 0.0 µg/mL), and multiple concentrations (1.3, 6.5, and 13.0 µg/mL) of alteplase (blue) and Brnoteplase (green). Box plots illustrate mean values (cross), median (line), interquartile range (box), and minimum/maximum values (whiskers). (**a**) Comparison of efficacy assessed by the time required to restore recanalization (if no recanalization achieved at the end of the 180 min experiment window, the value of '180' assigned). (**b**) Comparison of efficacy assessed by the recanalization frequency. (**c**) Comparison of efficacy assessed by the clot reduction at the end of the 180 min experiment window. (**d**) Comparison of efficacy assessed by the clot lysis rate over time.

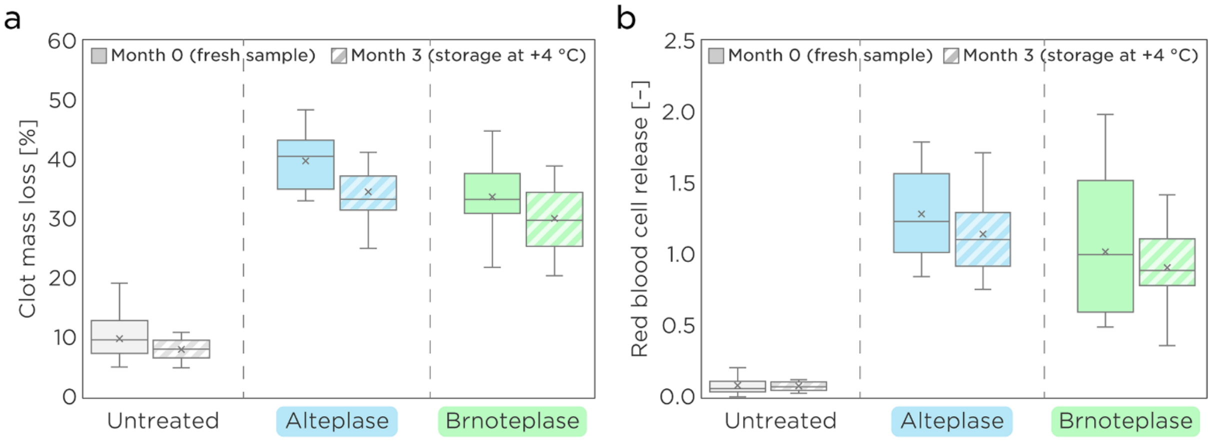

**Figure S12: *In vitro* analysis of clot lysis efficacy in the static model with semi-synthetic clots after long-term storage.** The comparison includes untreated group (grey), alteplase (blue), and Brnoteplase (green). The efficacy was measured at month 0 with freshly prepared samples (solid boxes) and again at month 3 after their incubation at +4 °C (hatched boxes). Box plots illustrate mean values (cross), median (line), interquartile range (box), and minimum/maximum values (whiskers). (**a**) Comparison of efficacy expressed as clot mass loss. (**b**) Comparison of efficacy expressed as red blood cell release.

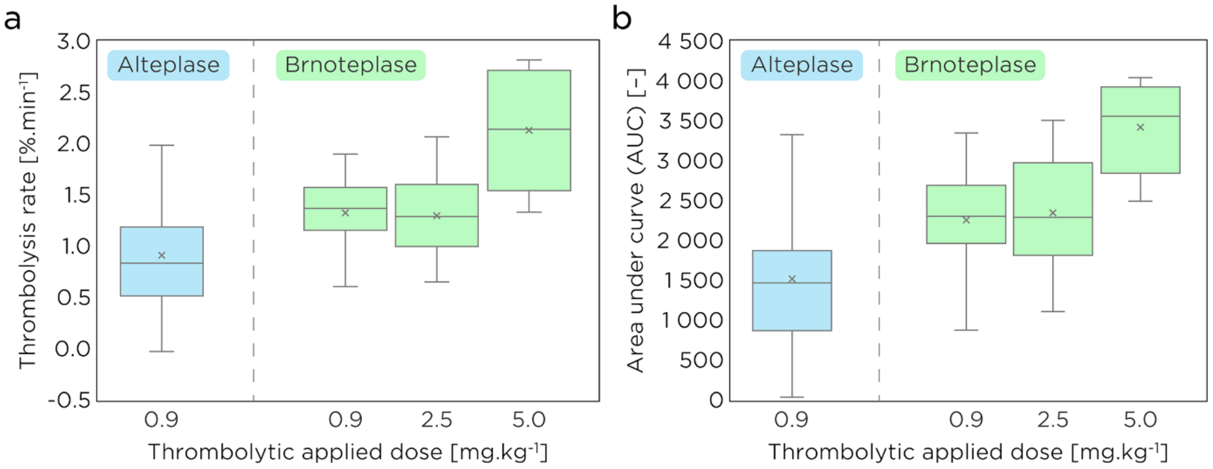

**Figure S13: Comparison of the *in vivo* thrombolytic effectivity in animal rat models at varying applied doses of tested thrombolytics.** The results were collected for alteplase (blue; administered as continuous rate infusion) at the clinically relevant dose of 0.9 mg/kg and for Brnoteplase (green; administered as bolus) at the dosses of 0.9, 2.5, and 5.0 mg/kg. (**a**) Thrombolytic effectivity expressed as lysis rate (% of the clot dissolved per minute). (**b**) Thrombolytic effectivity expressed as the area under the curve (AUC) of the lytic curves. Box plots show mean values (cross), median (line), interquartile range (box), and minimum/maximum values (whiskers).

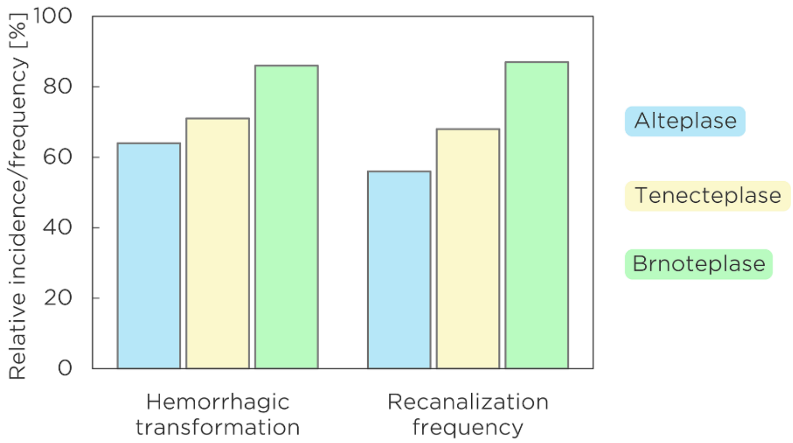

**Figure S14: Comparison of hemorrhagic transformation incidence and recanalization frequency observed *in vivo* in animal rat models.** The data were collected after the administration of alteplase (blue; 0.9 mg/kg; continuous rate infusion), tenecteplase (yellow; 0.25 mg/kg; bolus), and Brnoteplase (green; 2.5 mg/kg; bolus). The results show clear correlation between the overall hemorrhagic transformation incidence and recanalization frequency.

### Supplementary tables

**Table S1: Comparison of all key parameters derived from the biochemical characterization of thrombolytics.** The resulting data are presented as values ± standard error.

| Property | Key parameter | Alteplase | Brnoteplase | Alt01 | Alt 02 | Alt03 | AltAnc02 | AltEM03 | DesEM01 | DesEM02 | Desmoteaplse | Tenecteplase |
|---|---|---|---|---|---|---|---|---|---|---|---|---|
| Enzymatic activity | Activity (×$10^{-5}$) [$s^{-2}$] | 35.6 ± 7.8 | 2.2 ± 0.3 | 33.5 ± 3.1 | 24.2 ± 4.1 | 31.3 ± 3.7 | 26.8 ± 4.0 | 63.7 ± 2.9 | 45.3 ± 4.8 | 1.9 ± 0.2 | 1.9 ± 0.2 | 1.8 ± 0.2 |
| | Relative activity [-] | 1.00 ± 0.22 | 0.06 ± 0.01 | 0.94 ± 0.09 | 0.68 ± 0.12 | 0.88 ± 0.10 | 0.75 ± 0.11 | 1.79 ± 0.08 | 1.27 ± 0.14 | 0.05 ± 0.01 | 0.05 ± 0.01 | 0.05 ± 0.01 |
| | Fg-stimulated activity (×$10^{-5}$) [$s^{-2}$] | 725 ± 99 | 3.1 ± 1.1 | 652 ± 63 | 660 ± 210 | 551 ± 49 | 68 ± 14 | 120 ± 5 | 73 ± 21 | 1.0 ± 0.2 | 1.9 ± 0.3 | 2.9 ± 0.4 |
| | Relative Fg activity [-] | 1.00 ± 0.14 | 0.0043 ± 0.0015 | 0.90 ± 0.09 | 0.90 ± 0.29 | 0.76 ± 0.07 | 0.09 ± 0.02 | 0.17 ± 0.01 | 0.10 ± 0.03 | 0.0014 ± 0.0003 | 0.0027 ± 0.0004 | 0.0040 ± 0.0006 |
| | Fn-stimulated activity (×$10^{-5}$) [$s^{-2}$] | 9770 ± 510 | 3200 ± 280 | 6180 ± 200 | 6100 ± 280 | 8200 ± 1100 | 3780 ± 230 | 1360 ± 20 | 2010 ± 110 | 32 ± 4 | 5980 ± 360 | 3030 ± 120 |
| | Relative Fn activity [-] | 1.00 ± 0.05 | 0.33 ± 0.03 | 0.63 ± 0.02 | 0.62 ± 0.03 | 0.84 ± 0.11 | 0.39 ± 0.02 | 0.14 ± 0.01 | 0.21 ± 0.01 | 0.0033 ± 0.0004 | 0.61 ± 0.04 | 0.31 ± 0.01 |
| Stimulation and selectivity | Fg stimulation factor [-] | 20.4 ± 5.3 | 1.4 ± 0.5 | 19.5 ± 2.6 | 27.1 ± 9.7 | 17.6 ± 2.6 | 3.8 ± 1.0 | 2.8 ± 0.2 | 2.4 ± 0.7 | 0.8 ± 0.2 | 1.7 ± 0.3 | 1.6 ± 0.3 |
| | Fn stimulation factor [-] | 275 ± 62 | 1430 ± 220 | 185 ± 18 | 252 ± 45 | 260 ± 47 | 89 ± 14 | 14 ± 1 | 28 ± 3 | 10 ± 2 | 2210 ± 280 | 1650 ± 170 |
| | Selectivity [-] | 13.5 ± 2.0 | 1020 ± 360 | 9.5 ± 1.0 | 9.3 ± 3.0 | 14.8 ± 2.4 | 23.4 ± 4.9 | 4.8 ± 0.2 | 11.7 ± 3.4 | 13.5 ± 3.1 | 1320 ± 220 | 1050 ± 170 |
| Fibrin binding (penetrability) | $K_d$ [μM] | 0.57 ± 0.03 | 2.30 ± 0.20 | 3.90 ± 0.40 | 1.30 ± 0.10 | 1.60 ± 0.10 | 0.58 ± 0.07 | 0.59 ± 0.04 | 1.55 ± 0.12 | 1.14 ± 0.30 | 0.93 ± 0.02 | 0.78 ± 0.01 |
| | Relative Fn binding [-] | 1.00 ± 0.05 | 0.25 ± 0.02 | 0.15 ± 0.01 | 0.44 ± 0.03 | 0.36 ± 0.02 | 0.98 ± 0.12 | 0.97 ± 0.07 | 0.37 ± 0.03 | 0.50 ± 0.13 | 0.61 ± 0.01 | 0.73 ± 0.01 |
| | Maximal binding [%] | 97 ± 1 | 99 ± 1 | 100 ± 1 | 92 ± 2 | 97 ± 1 | 96 ± 4 | 86 ± 1 | 98 ± 3 | 91 ± 7 | 102 ± 1 | 107 ± 1 |
| Inhibition resistance | $IC_{50}$ [nM] | 61 ± 3 | 253 ± 17 | 147 ± 3 | 146 ± 5 | 158 ± 2 | 55 ± 1 | 27 ± 1 | 41 ± 10 | 150 ± 6 | 131 ± 18 | 278 ± 24 |
| | Relative inhibition resistance [-] | 1.00 ± 0.06 | 4.15 ± 0.28 | 2.40 ± 0.05 | 2.40 ± 0.08 | 2.60 ± 0.04 | 0.90 ± 0.08 | 0.44 ± 0.02 | 0.68 ± 0.17 | 2.46 ± 0.10 | 2.15 ± 0.29 | 4.56 ± 0.39 |
| Thermal stability | $T_m$ [°C] | 74.4 ± 0.1 | 61.2 ± 0.2 | 71.7 ± 0.1 | 73.9 ± 0.02 | 73.4 ± 0.01 | 87.2 ± 0.7 | 66.5 ± 1.4 | 71.3 ± 0.2 | 72.3 ± 0.1 | 69.2 ± 0.2 | 70.4 ± 0.5 |
| | $T_{onset}$ [°C] | 64.4 ± 0.1 | 46.8 ± 0.2 | 59.5 ± 8.2 | 91.9 ± 0.8 | 48.4 ± 16.1 | 48.9 ± 0.3 | 41.6 ± 0.9 | 53.1 ± 1.1 | 57.3 ± 8.4 | 48.1 ± 4.1 | 56.6 ± 5.1 |

**Table S2: Comparison of *in vitro* clot lysis efficacy for various thrombolytics in the static model with semi-synthetic clots.** The concentration of all the tested thrombolytics was 1.3 µg.mL$^{-1}$ and clots were incubated with the tested thrombolytic for 60 min at 37 °C. The comparison includes untreated group, alteplase, Alt01, Alt02, Alt03, and Brnoteplase.

| | Relative clot mass loss [%] | | | | | |
|---|---|---|---|---|---|---|
| | Untreated | Alteplase | Alt01 | Alt02 | Alt03 | Brnoteplase |
| mean | 10.7 | 41.2 | 47.1 | 42.3 | 39.5 | 31.0 |
| median | 10.3 | 41.6 | 41.3 | 40.5 | 41.3 | 30.8 |
| SD | 3.7 | 5.7 | 15.3 | 7.8 | 8.3 | 6.4 |
| minimum | 4.3 | 25.4 | 33.3 | 29.0 | 25.7 | 15.7 |
| maximum | 19.2 | 48.4 | 75.3 | 55.6 | 50.5 | 44.8 |
| count | 27 | 21 | 9 | 9 | 9 | 21 |
| | **Red blood cell release [-]** | | | | | |
| | Untreated | Alteplase | Alt01 | Alt02 | Alt03 | Brnoteplase |
| mean | 0.11 | 1.24 | 1.47 | 1.05 | 0.97 | 0.65 |
| median | 0.11 | 1.26 | 1.15 | 0.95 | 0.75 | 0.58 |
| SD | 0.09 | 0.37 | 0.80 | 0.40 | 0.51 | 0.32 |
| minimum | 0.01 | 0.39 | 0.71 | 0.64 | 0.36 | 0.13 |
| maximum | 0.28 | 1.79 | 2.83 | 1.93 | 1.84 | 1.52 |
| count | 9 | 21 | 9 | 9 | 9 | 21 |

**Table S3: Concentration dependence profiles of *in vitro* clot lysis efficacy in the static model with semi-synthetic clots.** Clots were incubated in human plasma with alteplase or Brnoteplase at different concentrations at 37 °C for 60 min. The lower and upper limits stand for confidence interval (95% CI, Gaussian distribution statistics).

**Relative clot mass loss [%]**

| | Control | Alteplase dose (μg.mL$^{-1}$) | | | | | | Brnoteplase dose (μg.mL$^{-1}$) | | | | | |
|---|---|---|---|---|---|---|---|---|---|---|---|---|---|
| | | 0.13 | 1.3 | 2.6 | 6.5 | 13 | 65 | 0.13 | 1.3 | 2.6 | 6.5 | 13 | 65 |
| mean | 10.5 | 17.1 | 41.2 | 44.8 | 42.2 | 38.7 | 31.1 | 9.6 | 31 | 34.2 | 39.7 | 44.1 | 41.8 |
| median | 10.2 | 15.3 | 41.6 | 44.9 | 45 | 36.8 | 30.3 | 8.3 | 30.8 | 32.9 | 39.9 | 41.8 | 41.4 |
| SD | 3.7 | 8 | 5.6 | 3.4 | 5.6 | 5.9 | 3.6 | 6.1 | 6.4 | 9.6 | 3 | 4.9 | 5.1 |
| lower 95% CI | 9.1 | 10.9 | 38.6 | 42.2 | 37.9 | 35.8 | 28.4 | 4.9 | 28.1 | 26.9 | 37.2 | 41.7 | 37.9 |
| upper 95% CI | 11.8 | 23.2 | 43.8 | 47.4 | 46.5 | 41.6 | 33.9 | 14.3 | 33.9 | 41.6 | 42.2 | 46.5 | 45.7 |
| count | 30 | 9 | 21 | 9 | 9 | 18 | 9 | 9 | 21 | 9 | 8 | 18 | 9 |

**Red blood cell release [-]**

| | Control | Alteplase dose (μg.mL$^{-1}$) | | | | | | Brnoteplase dose (μg.mL$^{-1}$) | | | | | |
|---|---|---|---|---|---|---|---|---|---|---|---|---|---|
| | | 0.13 | 1.3 | 2.6 | 6.5 | 13 | 65 | 0.13 | 1.3 | 2.6 | 6.5 | 13 | 65 |
| mean | 0.07 | 0.27 | 1.24 | 1.19 | 1.53 | 1.14 | 0.79 | 0.14 | 0.65 | 0.82 | 0.85 | 1.13 | 1.13 |
| median | 0.06 | 0.16 | 1.26 | 1.26 | 1.37 | 1.01 | 0.83 | 0.07 | 0.58 | 0.6 | 0.7 | 1.12 | 1.13 |
| SD | 0.06 | 0.23 | 0.37 | 0.35 | 0.44 | 0.46 | 0.17 | 0.2 | 0.32 | 0.52 | 0.43 | 0.4 | 0.29 |
| lower 95% CI | 0.05 | 0.09 | 1.07 | 0.92 | 1.2 | 0.91 | 0.66 | -0.01 | 0.5 | 0.41 | 0.49 | 0.93 | 0.91 |
| upper 95% CI | 0.09 | 0.44 | 1.4 | 1.46 | 1.87 | 1.37 | 0.92 | 0.29 | 0.8 | 1.22 | 1.21 | 1.33 | 1.35 |
| count | 30 | 9 | 21 | 9 | 9 | 18 | 9 | 9 | 21 | 9 | 8 | 18 | 9 |

**Table S4: Concentration dependence profiles of *in vitro* clot lysis efficacy in the static model with red blood cell dominant clots.** Clots were incubated in human plasma with alteplase or Brnoteplase at different concentrations at 37 °C for 60 min. The lower and upper limits stand for confidence interval (95% CI, Gaussian distribution statistics).

**Relative clot mass loss [%]**

| | Control | Alteplase dose (μg.mL$^{-1}$) | | | | | Brnoteplase dose (μg.mL$^{-1}$) | | | | |
|---|---|---|---|---|---|---|---|---|---|---|---|
| | | 0.13 | 1.3 | 2.6 | 6.5 | 13 | 0.13 | 1.3 | 2.6 | 6.5 | 13 |
| mean | 25.8 | 21.9 | 34.6 | 33.5 | 44.0 | 45.6 | 19.0 | 23.5 | 23.7 | 27.3 | 33.9 |
| median | 27.0 | 19.3 | 34.7 | 32.9 | 44.9 | 43.4 | 20.6 | 22.4 | 23.8 | 26.4 | 32.5 |
| SD | 5.7 | 7.0 | 8.6 | 7.2 | 9.7 | 8.4 | 5.5 | 4.7 | 7.9 | 5.5 | 8.5 |
| lower 95% CI | 23.0 | 16.5 | 27.4 | 27.9 | 36.6 | 39.2 | 14.7 | 19.9 | 17.6 | 23.1 | 26.8 |
| upper 95% CI | 28.6 | 27.2 | 41.8 | 39.0 | 51.5 | 52.1 | 23.2 | 27.1 | 29.7 | 31.6 | 41.1 |
| count | 18 | 9 | 8 | 9 | 9 | 9 | 9 | 9 | 9 | 9 | 8 |

**Red blood cell release [-]**

| | Control | Alteplase dose (μg.mL$^{-1}$) | | | | | Brnoteplase dose (μg.mL$^{-1}$) | | | | |
|---|---|---|---|---|---|---|---|---|---|---|---|
| | | 0.13 | 1.3 | 2.6 | 6.5 | 13 | 0.13 | 1.3 | 2.6 | 6.5 | 13 |
| mean | 0.60 | 0.78 | 2.00 | 2.09 | 3.32 | 3.59 | 0.58 | 0.68 | 0.70 | 1.13 | 1.70 |
| median | 0.49 | 0.97 | 1.88 | 2.21 | 2.68 | 3.72 | 0.45 | 0.68 | 0.58 | 1.13 | 1.43 |
| SD | 0.30 | 0.35 | 0.76 | 0.73 | 1.20 | 0.71 | 0.24 | 0.37 | 0.44 | 0.42 | 0.89 |
| lower 95% CI | 0.45 | 0.51 | 1.36 | 1.53 | 2.39 | 3.05 | 0.40 | 0.40 | 0.36 | 0.81 | 1.02 |
| upper 95% CI | 0.75 | 1.05 | 2.64 | 2.65 | 4.24 | 4.14 | 0.77 | 0.96 | 1.03 | 1.45 | 2.38 |
| count | 18 | 9 | 8 | 9 | 9 | 9 | 9 | 9 | 9 | 9 | 9 |

**Table S5: Comparison of *in vitro* clot lysis efficacy and concentration profiles in the flow model with semi-synthetic clots.** Clots were incubated in human plasma with alteplase or Brnoteplase at different concentrations at 37 °C for 180 min. The lower and upper limits stand for confidence interval (95% CI, Gaussian distribution statistics). N/A = not applicable

**Recanalization time [min]**

| | Control | Alteplase (µg.mL$^{-1}$) | | | Brnoteplase (µg.mL$^{-1}$) | | |
|---|---|---|---|---|---|---|---|
| | | 1.3 | 6.5 | 13 | 1.3 | 6.5 | 13 |
| mean | 180 | 144 | 176 | 180 | 173 | 146 | 158 |
| median | 180 | 147 | 180 | 180 | 175 | 152 | 180 |
| SD | 0 | 31 | 7 | 0 | 8 | 37 | 34 |
| lower 95% CI | N/A | 121 | 170 | N/A | 167 | 118 | 134 |
| upper 95% CI | N/A | 166 | 181 | N/A | 179 | 175 | 182 |
| count | 10 | 10 | 10 | 9 | 10 | 9 | 10 |

**Recanalization frequency [%]**

| | Control | Alteplase (µg.mL$^{-1}$) | | | Brnoteplase (µg.mL$^{-1}$) | | |
|---|---|---|---|---|---|---|---|
| | | 1.3 | 6.5 | 13 | 1.3 | 6.5 | 13 |
| mean | 0 | 100 | 30 | 0 | 50 | 60 | 40 |
| median | 0 | 100 | 0 | 0 | 50 | 50 | 50 |
| SD | 0 | 0 | 45 | 0 | 35 | 42 | 42 |
| lower 95% CI | N/A | N/A | -26 | N/A | 6 | 8 | -12 |
| upper 95% CI | N/A | N/A | 86 | N/A | 94 | 112 | 92 |
| count | 5 | 4 | 5 | 4 | 5 | 5 | 5 |

**Relative clot reduction [%]**

| | Control | Alteplase (µg.mL$^{-1}$) | | | Brnoteplase (µg.mL$^{-1}$) | | |
|---|---|---|---|---|---|---|---|
| | | 1.3 | 6.5 | 13 | 1.3 | 6.5 | 13 |
| mean | 1.4 | 100.0 | 51.4 | 15.9 | 64.3 | 69.8 | 55.7 |
| median | 0.0 | 100.0 | 35.7 | 14.3 | 71.4 | 100.0 | 35.7 |
| SD | 4.5 | 0.0 | 35.8 | 15.1 | 38.2 | 38.1 | 38.9 |
| lower 95% CI | -1.8 | N/A | 25.8 | 4.3 | 36.9 | 40.6 | 27.9 |
| upper 95% CI | 4.7 | N/A | 77.0 | 27.4 | 91.6 | 99.1 | 83.6 |
| count | 10 | 9 | 10 | 9 | 10 | 9 | 10 |

**Relative clot reduction rate [%.min$^{-1}$]**

| | Control | Alteplase (µg.mL$^{-1}$) | | | Brnoteplase (µg.mL$^{-1}$) | | |
|---|---|---|---|---|---|---|---|
| | | 1.3 | 6.5 | 13 | 1.3 | 6.5 | 13 |
| mean | 0.01 | 0.54 | 0.27 | 0.14 | 0.36 | 0.46 | 0.36 |
| median | 0.00 | 0.59 | 0.22 | 0.11 | 0.37 | 0.55 | 0.24 |
| SD | 0.02 | 0.18 | 0.15 | 0.13 | 0.19 | 0.27 | 0.26 |
| lower 95% CI | -0.01 | 0.41 | 0.17 | 0.05 | 0.23 | 0.25 | 0.17 |
| upper 95% CI | 0.02 | 0.67 | 0.38 | 0.23 | 0.50 | 0.67 | 0.54 |
| count | 10 | 10 | 10 | 10 | 10 | 9 | 10 |

**Red blood cell release [-]**

| | Control | Alteplase (µg.mL$^{-1}$) | | | Brnoteplase (µg.mL$^{-1}$) | | |
|---|---|---|---|---|---|---|---|
| | | 1.3 | 6.5 | 13 | 1.3 | 6.5 | 13 |
| mean | 0.02 | 0.16 | 0.08 | 0.06 | 0.15 | 0.19 | 0.15 |
| median | 0.02 | 0.16 | 0.09 | 0.05 | 0.16 | 0.21 | 0.14 |
| SD | 0.02 | 0.13 | 0.06 | 0.04 | 0.04 | 0.13 | 0.06 |
| lower 95% CI | 0.01 | 0.07 | 0.04 | 0.03 | 0.11 | 0.09 | 0.10 |
| upper 95% CI | 0.04 | 0.26 | 0.12 | 0.09 | 0.18 | 0.29 | 0.20 |
| count | 9 | 10 | 10 | 10 | 8 | 9 | 8 |

**Table S6: Comparison of long-term storage effect on *in vitro* clot lysis efficacy in the static model with semi-synthetic clots.** The concentration of all the tested thrombolytics was 1.3 μg.mL$^{-1}$ and clots were incubated with the tested thrombolytic for 60 min at 37 °C. 'Month 0' corresponds to fresh samples, 'month 3' corresponds to samples stored at +4 °C for three months.

| **Relative clot mass loss [%]** | | | | | | |
|---|---|---|---|---|---|---|
| | MONTH 0 | | | MONTH 3 | | |
| | Untreated | Alteplase | Brnoteplase | Untreated | Alteplase | Brnoteplase |
| mean | 9.8 | 39.8 | 33.6 | 8.0 | 34.5 | 30.1 |
| median | 9.6 | 40.6 | 33.3 | 8.0 | 33.3 | 29.8 |
| SD | 3.4 | 5.0 | 5.6 | 1.8 | 4.8 | 5.5 |
| minimum | 5.0 | 33.1 | 19.8 | 4.9 | 25.1 | 20.4 |
| maximum | 19.2 | 48.4 | 44.8 | 10.9 | 47.0 | 38.9 |
| count | 30 | 15 | 15 | 18 | 18 | 18 |

| **Red blood cell release [-]** | | | | | | |
|---|---|---|---|---|---|---|
| | MONTH 0 | | | MONTH 3 | | |
| | Untreated | Alteplase | Brnoteplase | Untreated | Alteplase | Brnoteplase |
| mean | 0.08 | 1.28 | 1.02 | 0.08 | 1.14 | 0.91 |
| median | 0.06 | 1.23 | 1.00 | 0.07 | 1.11 | 0.89 |
| SD | 0.06 | 0.31 | 0.46 | 0.03 | 0.28 | 0.27 |
| minimum | 0.00 | 0.85 | 0.49 | 0.03 | 0.76 | 0.36 |
| maximum | 0.23 | 1.79 | 1.99 | 0.12 | 1.72 | 1.42 |
| count | 28 | 14 | 15 | 17 | 17 | 18 |

**Table S7: Comparison of *in vitro* penetration capability of thrombolytic proteins through fibrin network.** The comparison includes fibrin non-interacting protein bovine serum albumin (BSA, control) and tested thrombolytics alteplase and Brnoteplase. The proteins were applied at concentration corresponding to clinically relevant dosing of alteplase indicated for patients with ischemic stroke (1.3 μg.mL$^{-1}$). Penetration rate was evaluated after 180 min at 37 °C. The lower and upper limits stand for confidence interval (95% CI, Gaussian distribution statistics).

| Normalized penetration rate [-] | | | | | | | | | | | | | | | |
|---|---|---|---|---|---|---|---|---|---|---|---|---|---|---|---|
| | BSA (control) | | | | | Alteplase | | | | | Brnoteplase | | | | |
| distance [mm] | 0.00 | 0.25 | 0.50 | 0.75 | 1.00 | 0.00 | 0.25 | 0.50 | 0.75 | 1.00 | 0.00 | 0.25 | 0.50 | 0.75 | 1.00 |
| mean | 1.0 | 1.0 | 0.9 | 0.9 | 0.9 | 0.5 | 0.4 | 0.3 | 0.2 | 0.2 | 0.6 | 0.5 | 0.4 | 0.4 | 0.4 |
| median | 1.0 | 1.0 | 0.9 | 0.9 | 0.9 | 0.4 | 0.4 | 0.4 | 0.2 | 0.2 | 0.5 | 0.5 | 0.4 | 0.4 | 0.4 |
| SD | 0.1 | 0.1 | 0.1 | 0.1 | 0.1 | 0.1 | 0.1 | 0.1 | 0.2 | 0.1 | 0.1 | 0.1 | 0.1 | 0.1 | 0.1 |
| lower 95% CI | 0.9 | 0.9 | 0.8 | 0.8 | 0.9 | 0.4 | 0.3 | 0.2 | 0.1 | 0.1 | 0.5 | 0.4 | 0.3 | 0.4 | 0.3 |
| upper 95% CI | 1.1 | 1.1 | 1.0 | 1.0 | 1.0 | 0.5 | 0.5 | 0.4 | 0.4 | 0.3 | 0.6 | 0.5 | 0.4 | 0.5 | 0.5 |
| count | 9 | 9 | 9 | 9 | 9 | 12 | 12 | 11 | 12 | 12 | 17 | 18 | 18 | 18 | 17 |

**Table S8: Comparison of *in vivo* thrombolytic effectivity in animal rat models.** The comparison includes untreated group (control), alteplase, Brnoteplase, and tenecteplase introduced at different doses (0.9, 2.5, 5.0, 0.25 mg/kg) and in different administration modes – continuous rate infusion (CRI) vs. bolus.

**Lysis rate [%.min$^{-1}$]**

| | Control | Alteplase (CRI) | Brnoteplase (bolus) | | | Tenecteplase (bolus) | Brnoteplase (CRI) |
|---|---|---|---|---|---|---|---|
| | | 0.9 mg/kg | 0.9 | 2.5 | 5.0 | 0.25 mg/kg | 0.9 mg/kg |
| mean | 0.41 | 0.91 | 1.33 | 1.30 | 2.13 | 1.08 | 0.41 |
| median | 0.39 | 0.84 | 1.37 | 1.29 | 2.14 | 0.97 | 0.41 |
| SD | 0.31 | 0.46 | 0.39 | 0.39 | 0.61 | 0.52 | 0.29 |
| lower 95% CI | 0.27 | 0.74 | 1.16 | 1.15 | 1.37 | 0.87 | 0.15 |
| upper 95% CI | 0.56 | 1.08 | 1.49 | 1.45 | 2.89 | 1.29 | 0.67 |
| count | 21 | 30 | 24 | 28 | 5 | 26 | 7 |

**Area under curve (AUC) [-]**

| | Control | Alteplase (CRI) | Brnoteplase (bolus) | | | Tenecteplase (bolus) | Brnoteplase (CRI) |
|---|---|---|---|---|---|---|---|
| | | 0.9 mg/kg | 0.9 mg/kg | 2.5 mg/kg | 5.0 mg/kg | 0.25 mg/kg | 0.9 mg/kg |
| mean | 660 | 1520 | 2260 | 2350 | 3420 | 2180 | 670 |
| median | 550 | 1500 | 2300 | 2290 | 3560 | 2000 | 720 |
| SD | 450 | 810 | 630 | 720 | 610 | 1060 | 520 |
| lower 95% CI | 460 | 1220 | 1990 | 2070 | 2670 | 1750 | 180 |
| upper 95% CI | 870 | 1820 | 2520 | 2630 | 4170 | 2610 | 1150 |
| count | 21 | 30 | 24 | 28 | 5 | 26 | 7 |

**Table S9: Comparison of *in vivo* thrombolytic safety and recanalization in animal rat models.** The comparison includes alteplase, Brnoteplase, and tenecteplase introduced at different doses (0.9, 2.5, and 0.25 mg/kg, respectively) and in different administration modes – continuous rate infusion (CRI) vs. bolus.

| Parameter | Alteplase (0.9 mg/kg; CRI) | Brnoteplase (2.5 mg/kg; bolus) | Tenecteplase (0.25 mg/kg; bolus) |
|---|---|---|---|
| Count (*n*) | 25 | 23 | 25 |
| Count included (*n* included) | 22 | 22 | 24 |
| HT count | 14 | 19 | 17 |
| Overall HT incidence [%] | 64 | 86 | 71 |
| HI1 proportion [%] | 43 | 60 | 24 |
| HI2 proportion [%] | 36 | 26 | 41 |
| PH1 proportion [%] | 21 | 15 | 35 |
| Recanalization frequency [%] | 56 | 87 | 68 |
| Hemispheric asymmetry [%] | 15 ± 11 | 11 ± 8 | 13 ± 5 |

**Table S10: Comparison of pharmacological marker levels in rat plasma samples collected at the end of *in vivo* thrombolytic effectivity experiments**. The comparison includes determination of the plasminogen/plasmin ratio and residual level of an active thrombolytic protein. The levels were determined for rat plasma samples collected after the treatment with alteplase or Brnoteplase at varying doses (0.9, 2.5, and 5.0 mg/kg).

| **Plasminogen/plasmin ratio [-]** | | | | | | |
|---|---|---|---|---|---|---|
| | Alteplase (continuous rate infusion) | | | Brnoteplase (bolus) | | |
| | 0.9 mg/kg | 2.5 mg/kg | 5.0 mg/kg | 0.9 mg/kg | 2.5 mg/kg | 5.0 mg/kg |
| Mean | 1970 | 1720 | 1550 | 3050 | 2150 | 2700 |
| Standard error | 510 | 340 | 200 | 280 | 380 | 130 |
| **Residual active thrombolytic level [mg.L$^{-1}$]** | | | | | | |
| | Alteplase (continuous rate infusion) | | | Brnoteplase (bolus) | | |
| | 0.9 mg/kg | 2.5 mg/kg | 5.0 mg/kg | 0.9 mg/kg | 2.5 mg/kg | 5.0 mg/kg |
| Mean | 0.0060 | 0.065 | 0.086 | 0.22 | 0.70 | 2.54 |
| Standard error | 0.0007 | 0.010 | 0.013 | 0.06 | 0.15 | 0.09 |

#### Supplementary notes

**Supplementary Note 1: Structure-based rational design of alteplase.** In the structure-based strategy, we were selecting lysines important for binding LRP1. Within the first four domains of tPA, which are responsible for binding LRP1, there are 12 lysines[1]. Four lysines are evolutionarily conserved: Lys212, 228, 240, and 274. Eight lysines are exposed on the protein surface, and therefore, can mediate the binding to LRP1: Lys10, 49, 124, 159, 162, 212, and 228. Lysines 212 and 228 are on the same side of the protein as Tyr67, which was confirmed to be important for binding LRP1. In the protein-protein docking using ClusPro and Rosetta, Lys10, 49, 212, 228, and 247 were in contact with the LRP1 complement-binding repeat. Lys49 was in contact with LRP1 using both used docking methods and is very close to Tyr67, making it likely to participate in LRP1 binding. Therefore, we measured the distances of other lysines to Lys49 during the molecular dynamics simulations. Lys10, Lys152, Lys212, Lys228, and Lys247 were in the distance of 21-29 Å required for LRP1 binding. Based on surface exposure, docking results, and correct distance in the molecular dynamics, we selected Lys10, Lys49, and Lys152. They were mutated to Glu, Ile, and Gln, respectively, based on prediction of thermodynamic stability. Moreover, mutation Tyr67Asn and Ser69Ala were introduced to prevent glycosylation, and therefore, further reduce LRP1 binding[2]. This resulted in mutant Alt01: Lys10Glu/Lys49Ile/Tyr67Asn/Ser69Ala/ Lys152Gln.

**Supplementary Note 2: Sequence-based rational design of alteplase.** In the sequence-based strategy, we have identified that desmoteplase, an enzyme derived from vampire bat saliva, which competitively inhibits LRP1 binding of alteplase, contains lysines at positions homologous to Lys82, 152, 159, and 162 of alteplase[3]. Since Lys152 was mutated already in Alt01, mutant Alt02 contains the Lys82Ala/Lys159Ser/Lys162Ile mutations. Tyr67Asn and Ser69Ala mutations were also introduced as in Alt01 to further prevent LRP1 binding[2].

**Supplementary Note 3: Ancestral sequence reconstruction of plasminogen activators.** FireProt$^{ASR}$ webserver was used to perform ancestral sequence reconstruction (ASR) of tPA-related proteins. The analysis has resulted in a phylogenetic tree with 150 extant sequences homologous to tPA and 149 sequences ancestral. We have selected nodes evolutionarily close to tPA to preserve thrombolytic function and specificity to human plasminogen and fibrin. These were nodes 242 and 235, which were named AltAnc01 and AltAnc02. AltAnc01 has 77 % sequential identity to tPA and is ancestral to tPA and 4 other extant sequences, which are all tissue-type plasminogen activators from different species. AltAnc01 also contains 2 gaps and 3 insertions, with aggregated length of 6 residues. AltAnc02 shows 71 % sequential identity to tPA and is ancestral to tPA and 11 other sequences, of which 10 are annotated as plasminogen activators and 1 is a hypothetical protein, likely also a plasminogen activator. Moreover, AltAnc02 also contains 2 gaps and 3 insertions, totaling 12 residues. Both ancestrals have preserved catalytic residues.

**Supplementary Note 4: Mining novel plasminogen activators from sequence databases.** EnzymeMiner webserver was used to search for new proteins using tPA as the query. The analysis yielded 4637 mined sequences, out of which we filtered 198 sequences with solubility equal to or higher than tPA, not containing transmembrane segments, and containing all domains and catalytic residues essential for tPA function. From these, we have manually selected AltEM01, AltEM02, and AltEM03, which are common bottlenose dolphin tissue plasminogen activator isoforms X1 and X2, and tissue plasminogen activator from Bolivian squirrel monkey, respectively. AltEM01, AltEM02, and AltEM03 share 78 %, 82 %, and 92 % sequence identity to tPA. Enzyme mining using desmoteplase as a query yielded 5379 sequences, from which 615 have met the solubility, functional domains, catalytic residues, and no transmembrane segments criteria. From these, we manually selected DesEM01 and DesEM02, which are the tissue-type plasminogen activator isoform X3 from Angolan Colobus, and the plasminogen activator from white-winged vampire bat. DesEM01 and DesEM02 share 68 % and 80 % sequence identity to desmoteplase, respectively.

**Supplementary Note 5: Static model analysis of thrombolysis on *in vitro* clots.** Treatment with both alteplase and Brnoteplase at concentration 1.3 $\mu g.mL^{-1}$ or higher provided greater clot mass loss ($p<0.001$) and red blood cell (RBC) release ($p<0.001$) compared to the untreated group for both clot types (semi-synthetic, RBC dominant). For semi-synthetic clots, alteplase provided the same clot mass loss as clinically relevant concentration (i.e., 1.3 $\mu g.mL^{-1}$) at concentrations 2.6, 6.5, and 13 $\mu g.mL^{-1}$ ($p=0.93$, $p>0.99$, $p=0.97$, respectively); and lower clot mass loss at concentrations 0.13 and 65 $\mu g.mL^{-1}$ ($p<0.001$, $p=0.002$, respectively). Brnoteplase provided the same clot mass loss as clinically relevant concentration (i.e., 1.3 $\mu g.mL^{-1}$) at concentration 2.6 $\mu g\ mL^{-1}$ ($p=0.96$); and higher clot mass loss at concentrations 6.5, 13, and 65 $\mu g.mL^{-1}$ ($p=0.02$, $p<0.001$, $p<0.001$, respectively); and lower clot mass loss at concentration 0.13 $\mu g.mL^{-1}$ ($p<0.001$). Alteplase provided higher clot mass loss compared to Brnoteplase at concentrations 1.3 and 2.6 $\mu g.mL^{-1}$ ($p<0.001$, $p=0.01$, respectively); the same clot mass loss at concentrations 0.13, 6.5, and 13 $\mu g.mL^{-1}$ ($p=0.24$, $p>0.99$, $p=0.20$, respectively); and lower clot mass loss at concentration 65 $\mu g.mL^{-1}$ ($p=0.009$). Collectively, the data indicated that alteplase-induced thrombolysis was already at saturation when using clinically relevant concentration, while such effect was shifted to higher concentrations with Brnoteplase. There was a gradual self-inhibition at higher concentrations with alteplase. Similar results were observed by RBC release. These results are shown in **Figure 3b**, **Figure S10a**, and **Table S3**. For RBC dominant clots, alteplase provided the same clot mass loss as clinically relevant concentration (i.e., 1.3 $\mu g.mL^{-1}$) at concentrations 2.6, 6.5, and 13 $\mu g.mL^{-1}$ ($p>0.99$, $p=0.24$, $p=0.09$, respectively); and lower clot mass loss at concentration 0.13 $\mu g.mL^{-1}$ ($p=0.02$). Brnoteplase provided the same clot mass loss as clinically relevant concentration (i.e., 1.3 $\mu g.mL^{-1}$) at concentrations 0.13, 2.6, 6.5, and 13 $\mu g.mL^{-1}$ ($p=0.95$, $p>0.99$, $p=0.98$, $p=0.13$, respectively). Alteplase provided the same clot mass loss as Brnoteplase at concentrations 0.13, 1.3, and 2.6 μg.mL$^{-1}$ ($p>0.99$, $p=0.08$, $p=0.16$, respectively); and higher clot mass loss at concentrations 6.5 and 13 μg.mL$^{-1}$ ($p<0.001$, $p=0.05$, respectively). In both cases there was a gradual increase of lytic efficacy dependent on concentration, though it was lower in Brnoteplase. Similar results were observed by RBC release. These results are shown in **Figure S10bc**, and **Table S4**.

**Supplementary Note 6: Flow model analysis of thrombolysis on *in vitro* clots.** Treatment with alteplase at concentration 1.3 μg.mL$^{-1}$ provided lower recanalization time ($p=0.005$) and greater clot reduction rate ($p<0.001$) and red blood cell (RBC) release ($p=0.006$) compared to the untreated group. At higher concentrations (6.5 and 13 μg.mL$^{-1}$), the efficacy was similar or marginally higher compared to the untreated group. Brnoteplase provided lower recanalization time and greater clot reduction rate and RBC release compared to the untreated group at all tested concentrations (1.3 μg.mL$^{-1}$: $p=0.03$, $p<0.001$, $p<0.001$; 6.5 μg.mL$^{-1}$: $p=0.03$, $p=0.001$, $p=0.006$; 13 μg.mL$^{-1}$: $p=0.07$, $p=0.002$, $p<0.001$; p-values for recanalization time, clot reduction rate, RBC release, respectively). Alteplase at concentration 1.3 μg.mL$^{-1}$ provided lower recanalization time, greater clot reduction rate, and similar RBC release compared to higher concentrations (vs. 6.5 μg.mL$^{-1}$: $p=0.05$, $p=0.05$, $p=0.31$; vs. 13 μg.mL$^{-1}$: $p=0.03$, $p<0.001$, $p=0.10$; p-values for recanalization time, clot reduction rate, RBC release, respectively). Brnoteplase at concentration 1.3 μg.mL$^{-1}$ provided the same recanalization time, clot reduction rate, and RBC release as at higher concentrations (vs. 6.5 μg.mL$^{-1}$: $p=0.19$, $p=0.89$, $p=0.93$; vs. 13 μg.mL$^{-1}$: $p=0.74$, $p>0.99$, $p>0.99$; p-values for recanalization time, clot reduction rate, RBC release, respectively). Both alteplase and Brnoteplase applied at the same concentration (1.3; 6.5; 13 μg.mL$^{-1}$) provided similar recanalization time (1.3 μg.mL$^{-1}$: $p=0.10$; 6.5.μg mL$^{-1}$: $p=0.11$; 13 μg.mL$^{-1}$: $p=0.38$), clot reduction rate (1.3 μg.mL$^{-1}$: $p=0.35$; 6.5 μg.mL$^{-1}$: $p=0.33$; 13 μg.mL$^{-1}$: $p=0.16$), and RBC release (1.3 μg.mL$^{-1}$: $p>0.99$; 6.5 μg.mL$^{-1}$: $p=0.11$; 13 μg.mL$^{-1}$: $p=0.29$). These results are shown in **Figure 3c**, **Figure S11**, and **Table S5**.

### Supplementary methods

#### Computer-aided selection of mutations alleviating binding of tPA to LRP1

**Acquisition and preparation of structures.** The three-dimensional structure of tPA determined using small-angle X-ray scattering was kindly provided by Dr. Ashish upon request[4]. Missing side chains, heavy atoms, and hydrogens were added, and orientations of Asn, Gln, and His side chains were optimized using the $H^{++}$ server at 0.1 mM salinity and pH 7.5[5,6]. The X-ray crystallography structure of the LRP1 complement-binding repeat was obtained from the Protein Data Bank (PDB ID: 2FCW)[7-9].

**Assessment of conservation and correlation of lysine residues.** The structure of tPA was submitted to the Hotspot Wizard 3.0 web server[10]. Lysine residues were assessed for their conservation, solvent-accessible surface area, and frequencies of other amino acids at their positions.

**Optimizing tPA structure using Rosetta FastRelax.** Backbone dihedral angles in disallowed regions and steric clashes in the structure of tPA were resolved using Rosetta 3.9 RosettaScripts Fastrelax protocol using the scoring function ref2015[11,12]. Chi2 dihedral angles were sampled only for Phe, Tyr, and His residues. The Fastrelax protocol was run with coordinate constraints on non-backbone heavy atoms created by the BOUNDED function. The constraints had a tolerance of 0.01 Å and a standard deviation of 0.5 Å. The relaxed structure was deprotonated as described previously using the $H^{++}$ server.

***In silico* glycosylation.** Oligosaccharides corresponding to the most common glycoform of type 2 tPA were constructed using the GLYCAM carbohydrate builder and positioned to the N-linking Asn residues using PyMOL[13].

**Parametrization.** Input topologies and coordinates were prepared using the Tleap module of AMBER 16[14]. The protein and glycan parts of the system were parametrized using the ff14SB[15] and GLYCAM_06j-1[16] force fields, respectively. The system was solvated in a box of water molecules so that all protein atoms were at least 20 Å from the box surface, so the system, including carbohydrate chains, is contained within the periodic box. The OPC3 water model[17] was used. The system was neutralized using 4 $Na^{+}$ ions for equilibration. The number of ions added for the production simulations was determined using the average volume in the last stage of equilibration simulation to achieve a final salinity of 0.154 M as in human blood[18]. The masses of the hydrogen atom-containing groups from the output topology files were repartitioned using AmberTools17 ParmED command HmassRepartition for all simulations[19].

**Molecular dynamics simulations.** Energy minimization and MD simulations were performed by using the PMEMD.CUDA module of AMBER14[20,21]. Initially, the system was minimized by 10 steps of steepest descent followed by 9990 steps of conjugate gradient with 500 $kcal.mol^{-1}.Å^{-2}$ restraints on all atoms of the glycoprotein. The system was further minimized in four more rounds, each consisting of 2500 steps of steepest descent and 7500 steps of conjugate gradient minimization with decreasing harmonic restraints. The restraints were applied as follows: 500, 125, and 25 kcal.mol$^{-1}$Å$^{-2}$ on the backbone atoms of the glycoprotein. Finally, the system was minimized with 5000 steps of steepest descent and 15000 steps of conjugate gradient minimization without restraints. The subsequent MD simulations employed periodic boundary conditions, the Particle Mesh Ewald method to treat electrostatic interactions, a 10 Å cut-off for nonbonded interactions, the SHAKE algorithm to fix all bonds containing hydrogens, and a 4 fs time step[22-24]. Equilibration simulations consisted of two steps: 40 ps of gradual heating from 0 to 310 K using the Langevin thermostat with a collision frequency of 1.0 ps$^{-1}$, constant pressure of 1.0 bar using the Berendsen barostat with a pressure relaxation time of 1.0 ps$^{-1}$, and harmonic restraints of 200.0 kcal.mol$^{-1}$Å$^{-2}$ on the positions of all glycoprotein atoms[25,26]. Then, the system was further equilibrated during 8,800 ps at 310 K using the Langevin thermostat with a collision frequency of 1.0 ps$^{-1}$ and constant pressure of 1.0 bar using the Berendsen barostat with a pressure coupling constant of 1.0 ps $^{-1}$ in 11 rounds of 800 ps each with decreasing harmonic restraints. The restraints were applied as follows: 125.0, 100.0, 75.0, 50.0, 25, 15.0, 10.0, 5.0, 1.0, 0.5, and 0.0 kcal.mol$^{-1}$Å$^{-2}$ on the backbone atoms of the glycoprotein. After equilibration, 20 ns long production MD simulations were run using the same settings as the last equilibration step. Coordinates were saved at intervals of 4 ps. The simulation was run in 3 replicas, producing 60 ns of aggregated simulation time.

**Analysis of molecular dynamics simulations.** The MD simulations resulting trajectories were analyzed using the CPPTRAJ module of AMBER14[27]. Distances of the C-alpha atoms of all lysines on the first four domains of tPA to each other during the course of the simulation were plotted. The simulations were also visualized in Caver Analyst 2.0 beta[28].

**Protein-protein docking using Cluspro.** tPA without carbohydrate chains and the LRP1 binding repeat were docked using the ClusPro web server[29]. The default settings were used, except for selecting the "others" mode of energy terms weights.

**Protein-protein docking using RosettaDock.** tPA without carbohydrate chains and the LRP1 binding repeat were docked by the Rosetta 2018.33 RosettaDock program using scoring function ref2015[11,30,31]. Global docking of LRP1 binding repeat was performed (flags: -spin, -randomize1, -randomize2, -dock_pert 3.0 8). The bound conformations were optimized with side-chain repacking and minimization (flags: -docking:sc_min true, -packing:repack_only.)

**Clustering of docked structures.** RosettaDock results were clustered using the Calibur application. We used all four clustering strategies of Calibur[32] and a maximum RMSD of structures in cluster of either 15.0, 10.0, or 5.0 Å. All cluster centers were visually inspected in PyMOL.

#### Sequence-based strategy

The sequences of tPA and desmoteplase were aligned using the ClustalΩ algorithm[33] on its web server with default settings. Lysines on the first four domains of tPA were compared with homologous positions in the desmoteplase sequence.

#### Selection of residues to mutate into

**Structure preparation.** From the molecular dynamics simulations trajectories, three representative structures were obtained by K-means clustering the 10 most populated clusters using the CPPTRAJ module of AMBER17 with the cluster population percentages reported. The clusters from the three replicas did not differ significantly from each other, so the representative structure from the most populated cluster from the first replica was chosen as the starting structure for ΔΔG calculations.

**Screening of substitution mutations using Rosetta ddg_monomer, Protocols 3 and 16.** ΔΔG of mutated positions designated for mutation was calculated using ddg_monomer protocols 3 and 16[34]. Rosetta 2018.33 and the Talaris2014 score function were used. (flags -corrections::restore_talaris_behavior, -score:weights talaris2014, -ddg::minimization_scorefunction talaris2014.wts, -ddg::minimization_patch)[11]. The renumbered structure of tPA was minimized with harmonic constraints placed on the C-alpha atoms in the crystal structure using the minimize_with_cst.default.linuxgccrelease program. The soft-repulsive design energy function (--soft_rep_design weights) was used for repacking side chains (--sc_min_only false). Optimization was performed on the whole protein without distance restriction (--local_opt_only false). The previously created constraint files were used during backbone minimization (--min_cst true). Optimization was performed on an 8 Å wide shell around the point mutation (--local_opt_only true), and checkpointing was suppressed (--suppress_checkpointing). The mean energy from 50 iterations was used as the final parameter describing the stability effects of single-point mutations. These options constitute the original protocol 3 described previously[34]. Substitution mutations with ΔΔG < 0 were also modeled by protocol 16. According to the original ddg monomer paper, the score function Talaris2014 was used, with the flag -corrections::restore_talaris_behavior as in the original ddg_monomer approach. Talaris2014 score function was used. The soft-repulsive design energy function (--soft_rep_design) was used for repacking side chains (--sc_min_only false). Optimization without distance restriction (--local_opt_only false). The previously created constraint files were used during backbone minimization (--min_cst true). Three rounds of optimization with increasing weight on the repulsive term (--ramp_repulsive true) were applied. This constitutes protocol 16. The final ΔΔG was calculated as the difference of the mean of the 20 best wild-type structures and the 20 best mutant structures.

#### Combining beneficial tPA mutations from the literature

Data about the mechanism of effectivity, side effects, and biological half-life caused by tPA and previous protein engineering studies were studied using literature research. We have chosen a combination of favorable mutations for biochemical characterization. Findings have been published in our review article[1].

#### Ancestral sequence reconstruction of tPA-related proteins

**Reconstruction of ancestral sequences.** The sequence of tPA, including the signal peptide, was submitted to the FireProt$^{ASR}$ web server[35]. The calculation was done using default settings.

**Assessment of differences of ancestral sequences and tPA.** Multiple sequence alignment of tPA and selected ancestral sequences was done using Clustal-Ω. Possible differences in key sites on tPA were visually inspected in JalView 2[36].

#### Mining novel enzymes from sequence databases

**Enzyme mining.** Novel thrombolytic enzymes were searched in protein sequence databases using EnzymeMiner[37]. Two separate runs were performed using the sequences of tPA (Uniprot ID P00750) and desmoteplase (P98119)[38]. The signal peptide was included in both sequences.

**Selecting novel enzymes for biochemical characterization.** Proteins for biochemical characterization were selected using negative and positive selection. Negative selection consisted of excluding proteins with a transmembrane region and mutation of catalytic residues. Positive selection was selecting proteins that are predicted to be more soluble than the protein used as a query for database search (0.647 for tPA, 0.527 for desmoteplase) and contain key domains for plasminogen activation and fibrin binding (trypsin-like serine protease PF00089, kringle PF00051, fibronectin-like finger PF00039)[39]. After automatically filtering for these characteristics, proteins with interesting properties were selected manually.

#### References

1. Mican J, Toul M, Bednar D, Damborsky J. Structural biology and protein engineering of thrombolytics. *Comput Struct Biotechnol J*. 2019;17:917-938. doi:10.1016/j.csbj.2019.06.023

2. Bassel-Duby R, Jiang NY, Bittick T, et al. Tyrosine 67 in the epidermal growth factor-like domain of tissue-type plasminogen activator is important for clearance by a specific hepatic receptor. *J Biol Chem*. 1992;267(14):9668-9677.

3. López-Atalaya JP, Roussel BD, Ali C, et al. Recombinant *Desmodus rotundus* salivary plasminogen activator crosses the blood-brain barrier through a low-density lipoprotein receptor-related protein-dependent mechanism without exerting neurotoxic effects. *Stroke*. 2007;38(3):1036-1043. doi:10.1161/01.STR.0000258100.04923.84

4. Rathore YS, Rehan M, Pandey K, Sahni G, Ashish. First structural model of full-length human tissue-plasminogen activator: A SAXS data-based modeling study. *J Phys Chem B*. 2012;116(1):496-502. doi:10.1021/jp207243n

5. Gordon JC, Myers JB, Folta T, Shoja V, Heath LS, Onufriev A. H++: A server for estimating pKas and adding missing hydrogens to macromolecules. *Nucleic Acids Res*. 2005;33:W368-371. doi:10.1093/nar/gki464

6. Anandakrishnan R, Aguilar B, Onufriev AV. H++ 3.0: Automating pK prediction and the preparation of biomolecular structures for atomistic molecular modeling and simulations. *Nucleic Acids Res*. 2012;40:W537-541. doi:10.1093/nar/gks375

7. Berman H, Henrick K, Nakamura H. Announcing the worldwide Protein Data Bank. *Nat Struct Mol Biol*. 2003;10(12):980-980. doi:10.1038/nsb1203-980

8. Berman HM, Westbrook J, Feng Z, et al. The Protein Data Bank. *Nucleic Acids Res*. 2000;28(1):235-242. doi:10.1093/nar/28.1.235

9. Fisher C, Beglova N, Blacklow SC. Structure of an LDLR-RAP complex reveals a general mode for ligand recognition by lipoprotein receptors. *Mol Cell*. 2006;22(2):277-283. doi:10.1016/j.molcel.2006.02.021

10. Sumbalova L, Stourac J, Martinek T, Bednar D, Damborsky J. HotSpot Wizard 3.0: Web server for automated design of mutations and smart libraries based on sequence input information. *Nucleic Acids Res*. 2018;46(W1):W356-W362. doi:10.1093/nar/gky417

11. Alford RF, Leaver-Fay A, Jeliazkov JR, et al. The Rosetta all-atom energy function for macromolecular modeling and design. *J Chem Theory Comput*. 2017;13(6):3031-3048. doi:10.1021/acs.jctc.7b00125

12. Fleishman SJ, Leaver-Fay A, Corn JE, et al. RosettaScripts: A scripting language interface to the Rosetta macromolecular modeling suite. *PLOS ONE*. 2011;6(6):e20161. doi:10.1371/journal.pone.0020161

13. Spellman MW, Basa LJ, Leonard CK, et al. Carbohydrate structures of human tissue plasminogen activator expressed in Chinese Hamster Ovary cells. *J Biol Chem*. 1989;264(24):14100-14111. doi:10.1016/S0021-9258(18)71649-9

14. Case DA, Cheatham III TE, Darden T, et al. The Amber biomolecular simulation programs. *J Comput Chem*. 2005;26(16):1668-1688. doi:10.1002/jcc.20290

15. Maier JA, Martinez C, Kasavajhala K, Wickstrom L, Hauser KE, Simmerling C. ff14SB: Improving the accuracy of protein side chain and backbone parameters from ff99SB. *J Chem Theory Comput*. 2015;11(8):3696-3713. doi:10.1021/acs.jctc.5b00255

16. Kirschner KN, Yongye AB, Tschampel SM, et al. GLYCAM06: A generalizable biomolecular force field. Carbohydrates. *J Comput Chem*. 2008;29(4):622-655. doi:10.1002/jcc.20820

17. Izadi S, Onufriev AV. Accuracy limit of rigid 3-point water models. *J Chem Phys*. 2016;145(7):074501. doi:10.1063/1.4960175

18. Awad S, Allison SP, Lobo DN. The history of 0.9% saline. *Clin Nutr*. 2008;27(2):179-188. doi:10.1016/j.clnu.2008.01.008

19. Hopkins CW, Le Grand S, Walker RC, Roitberg AE. Long-time-step molecular dynamics through hydrogen mass repartitioning. *J Chem Theory Comput*. 2015;11(4). doi:10.1021/ct5010406

20. Salomon-Ferrer R, Götz AW, Poole D, Le Grand S, Walker RC. Routine microsecond molecular dynamics simulations with AMBER on GPUs. 2. Explicit solvent particle mesh Ewald. *J Chem Theory Comput*. 2013;9(9):3878-3888. doi:10.1021/ct400314y

21. Götz AW, Williamson MJ, Xu D, Poole D, Le Grand S, Walker RC. Routine microsecond molecular dynamics simulations with AMBER on GPUs. 1. Generalized born. *J Chem Theory Comput*. 2012;8(5):1542-1555. doi:10.1021/ct200909j

22. Darden T, York D, Pedersen L. Particle mesh Ewald: An N·log(N) method for Ewald sums in large systems. *J Chem Phys*. 1993;98(12):10089-10092. doi:10.1063/1.464397

23. Ryckaert JP, Ciccotti G, Berendsen HJC. Numerical integration of the cartesian equations of motion of a system with constraints: Molecular dynamics of *n*-alkanes. *J Comput Phys*. 1977;23(3):327-341. doi:10.1016/0021-9991(77)90098-5

24. Miyamoto S, Kollman PA. Settle: An analytical version of the SHAKE and RATTLE algorithm for rigid water models. *J Comput Chem*. 1992;13(8):952-962. doi:10.1002/jcc.540130805

25. Berendsen HJC, Postma JPM, van Gunsteren WF, DiNola A, Haak JR. Molecular dynamics with coupling to an external bath. *J Chem Phys*. 1984;81(8):3684-3690. doi:10.1063/1.448118

26. Loncharich RJ, Brooks BR, Pastor RW. Langevin dynamics of peptides: The frictional dependence of isomerization rates of N-acetylalanyl-N'-methylamide. *Biopolymers*. 1992;32(5):523-535. doi:10.1002/bip.360320508

27. Roe DR, Cheatham TEI. PTRAJ and CPPTRAJ: Software for processing and analysis of molecular dynamics trajectory data. *J Chem Theory Comput*. 2013;9(7):3084-3095. doi:10.1021/ct400341p

28. Jurcik A, Bednar D, Byska J, et al. CAVER Analyst 2.0: Analysis and visualization of channels and tunnels in protein structures and molecular dynamics trajectories. *Bioinforma Oxf Engl*. 2018;34(20):3586-3588. doi:10.1093/bioinformatics/bty386

29. Kozakov D, Hall DR, Xia B, et al. The ClusPro web server for protein-protein docking. *Nat Protoc*. 2017;12(2):255-278. doi:10.1038/nprot.2016.169

30. Chaudhury S, Berrondo M, Weitzner BD, Muthu P, Bergman H, Gray JJ. Benchmarking and analysis of protein docking performance in Rosetta v3.2. *PLOS ONE*. 2011;6(8):e22477. doi:10.1371/journal.pone.0022477

31. Park H, Bradley P, Greisen Jr P, et al. Simultaneous optimization of biomolecular energy functions on features from small molecules and macromolecules. *J Chem Theory Comput*. 2016;12(12):6201-6212. doi:10.1021/acs.jctc.6b00819

32. Li SC, Ng YK. Calibur: A tool for clustering large numbers of protein decoys. *BMC Bioinformatics*. 2010;11(1):25. doi:10.1186/1471-2105-11-25

33. Sievers F, Wilm A, Dineen D, et al. Fast, scalable generation of high-quality protein multiple sequence alignments using Clustal Omega. *Mol Syst Biol*. 2011;7(1):MSB201175. doi:10.1038/msb.2011.75

34. Kellogg EH, Leaver-Fay A, Baker D. Role of conformational sampling in computing mutation-induced changes in protein structure and stability. *Proteins Struct Funct Bioinforma*. 2011;79(3):830-838. doi:10.1002/prot.22921

35. Musil M, Khan RT, Beier A, et al. FireProt$^{ASR}$: A web server for fully automated ancestral sequence reconstruction. *Brief Bioinform*. 2021;22(4):bbaa337. doi:10.1093/bib/bbaa337

36. Waterhouse AM, Procter JB, Martin DMA, Clamp M, Barton GJ. Jalview version 2—A multiple sequence alignment editor and analysis workbench. *Bioinformatics*. 2009;25(9):1189-1191. doi:10.1093/bioinformatics/btp033

37. Hon J, Borko S, Stourac J, et al. EnzymeMiner: Automated mining of soluble enzymes with diverse structures, catalytic properties and stabilities. *Nucleic Acids Res*. 2020;48(W1):W104-W109. doi:10.1093/nar/gkaa372

38. The UniProt Consortium. UniProt: The Universal Protein Knowledgebase in 2023. *Nucleic Acids Res*. 2023;51(D1):D523-D531. doi:10.1093/nar/gkac1052

39. Hon J, Marusiak M, Martinek T, et al. SoluProt: Prediction of soluble protein expression in *Escherichia coli*. *Bioinformatics*. 2021;37(1):23-28. doi:10.1093/bioinformatics/btaa1102
